# Spatiotemporal dynamics of sensory neuron and Merkel-cell remodeling are decoupled during epidermal homeostasis

**DOI:** 10.1101/2023.02.14.528558

**Authors:** Rachel C. Clary, Blair A. Jenkins, Ellen A. Lumpkin

## Abstract

As the juncture between the body and environment, epithelia are both protective barriers and sensory interfaces that continually renew. To determine whether sensory neurons remodel to maintain homeostasis, we used *in vivo* two-photon imaging of somatosensory axons innervating Merkel cells in adult mouse skin. These touch receptors were highly plastic: 63% of Merkel cells and 89% of branches appeared, disappeared, grew, regressed and/or relocated over a month. Interestingly, Merkel-cell plasticity was synchronized across arbors during rapid epithelial turnover. When Merkel cells remodeled, the degree of plasticity between Merkel-cell clusters and their axons was well correlated. Moreover, branches were stabilized by Merkel-cell contacts. These findings highlight the role of epithelial-neural crosstalk in homeostatic remodeling. Conversely, axons were also dynamic when Merkel cells were stable, indicating that intrinsic neural mechanisms drive branch plasticity. Two terminal morphologies innervated Merkel cells: transient swellings called boutons, and stable cups termed kylikes. In *Atoh1* knockout mice that lack Merkel cells, axons showed higher complexity than control mice, with exuberant branching and no kylikes. Thus, Merkel cells limit axonal branching and promote branch maturation. Together, these results reveal a previously unsuspected high degree of plasticity in somatosensory axons that is biased, but not solely dictated, by plasticity of target epithelial cells. This system provides a platform to identify intrinsic and extrinsic mechanisms that govern axonal patterning in epithelial homeostasis.

## Introduction

Sensory epithelia perform two vital functions: they serve as water-tight protective barriers to guard our tissues against environmental insults, and as sensory interfaces that encode the features of our world. Epithelia are densely innervated by axons, which are tasked with reliably encoding sensory information; however, to maintain a healthy barrier, epithelial cells continually renew (Alberts, 2002; Zimmerman et al., 2014; Whitehead et al., 1985; Graziadei & Metcalf, 1971; Graziadei & Monti-Gaziadei, 1979). This raises the question of whether sensory neurons likewise remodel during normal tissue homeostasis. Other types of axons throughout the nervous system show various degrees of structural plasticity during adulthood – at one extreme, motor axons innervating the neuromuscular junction are completely stable after a period of developmental pruning, whereas axons and dendrites in the central nervous system show structural changes that reflect alterations in synaptic function in many contexts (Bareyre et al., 2011; Botchkarev et al., 1997; Cheng et al., 2010; De Paola et al., 2006; Gangadharan et al., 2022; Lichtman et al., 1987; Luo et al., 2016; Takahashi et al., 2019; Trachtenberg et al., 2002; Balice-Gordon & Lichtman, 1993; Waller, 1850). By contrast, little is known about the degree to which sensory axons remodel during epithelial turnover.

The epidermis, which is skin’s epithelial layer, is innervated by different classes of somatosensory neurons that trigger our senses of touch, itch and pain. Depending on their sensory modality, these sensory afferents form unique axonal arbors at distinct, epithelial-derived target structures. These sensory domains renew on different timescales. For example, hair follicles, which are innervated by touch-sensitive axons, regenerate over weeks to months. By contrast, interfollicular epidermal cells, which are innervated by pruritoceptive and nociceptive neurons, renew every four weeks (Zimmerman et al., 2014; Halprin, 1972; Blanpain & Fuchs, 2009). Touch domes, which are raised structures adjacent to some hair follicles, have specialized epithelial cells, vasculature and dermal cell types distinct from those in hair follicles or the interfollicular epidermis (Iggo & Muir, 1969; Smith, 1977). These zones are home to Merkel cells, which are mechanosensory epithelial cells that are innervated by low-threshold mechanosensory axons. Together, these two components form a touch-receptor complex that encodes both gentle pressure and object features (Iggo & Muir, 1969; Johnson, 2001; Maksimovic et al., 2014, Maricich et al., 2012; Wellnitz et al., 2010; Woodbury & Koerber, 2007). Several studies have investigated the maintenance of Merkel cells in adult epidermis (Wright et al., 2017; Doucet et al., 2013), but the stability of the axons innervating these presynaptic epithelial cells has not been investigated. Moreover, this two-part touch receptor enables investigation into whether neuronal patterning and remodeling depends on Merkel cells and vice versa.

To address these open questions, we developed a platform to perform quantitative, longitudinal imaging of identified Merkel-cell afferents and Merkel cells for up to a month. Because Merkel cells form presynaptic-like active zones with sensory axons, we hypothesized that Merkel-cell turnover would stimulate remodeling of their associated axonal terminals. To our surprise, we instead found that sensory axon terminals are highly dynamic even when Merkel cells are stable.

## Results

To visualize structural plasticity in mechanosensory axons during skin homeostasis, we developed a preparation to perform *in vivo* time-lapse imaging of molecularly identified sensory neurons and Merkel-cell clusters in adult mouse skin. Whole-mount immunohistochemistry was used to verify transgenic markers of axon terminals and Merkel cells with established antibody labels of these cell types (**Figure 1**). In skin, the transcription factor *Atoh1 is* selectively expressed in Merkel cells (Lumpkin et al., 2003); therefore, we used an Atoh1-GFP fusion protein (Atoh1-A1GFP mouse line) as a marker of Merkel cells. Keratin 8-positive Merkel cells (K8; Kim & Holbrook, 1995) co-expressed Atoh1-GFP (**Figure 1A**; Rose et al., 2009, Marshall et al., 2016). Likewise, Merkel-cell innervating sensory axons that were immunoreactive for myelinated neuron marker neurofilament heavy (NFH) also expressed tdTomato driven by the endogenous TrkC promoter (TrkC-tdTomato; **Figure 1B–C**), which labels circumferential endings at the base of hair follicles and a small percentage of free nerve endings in addition to Merkel-cell afferents (Bai et al., 2015). We noted that TrkC-tdTomato fills a larger extent of axon terminals than NFH, allowing for visualization of fine branches of the Merkel cell afferent that are not labeled by NFH (**Figure 1C**). Next, we confirmed that number of Merkel cells and terminal branches per touch dome were comparable in the cranium, a common site for *in vivo* imaging, and back skin, which was a focus of previous studies (**Figure 1D–G;** Moll et al., 1996; Nakafusa et al., 2006; Marshall et al., 2016). These findings validate the use of the transgenic markers TrkC-tdTomato and Atoh1-A1GFP to visualize mechanosensory axons and presynaptic epithelial cells on the crown of the head to study structural plasticity during tissue homeostasis.

**Figure 1.**
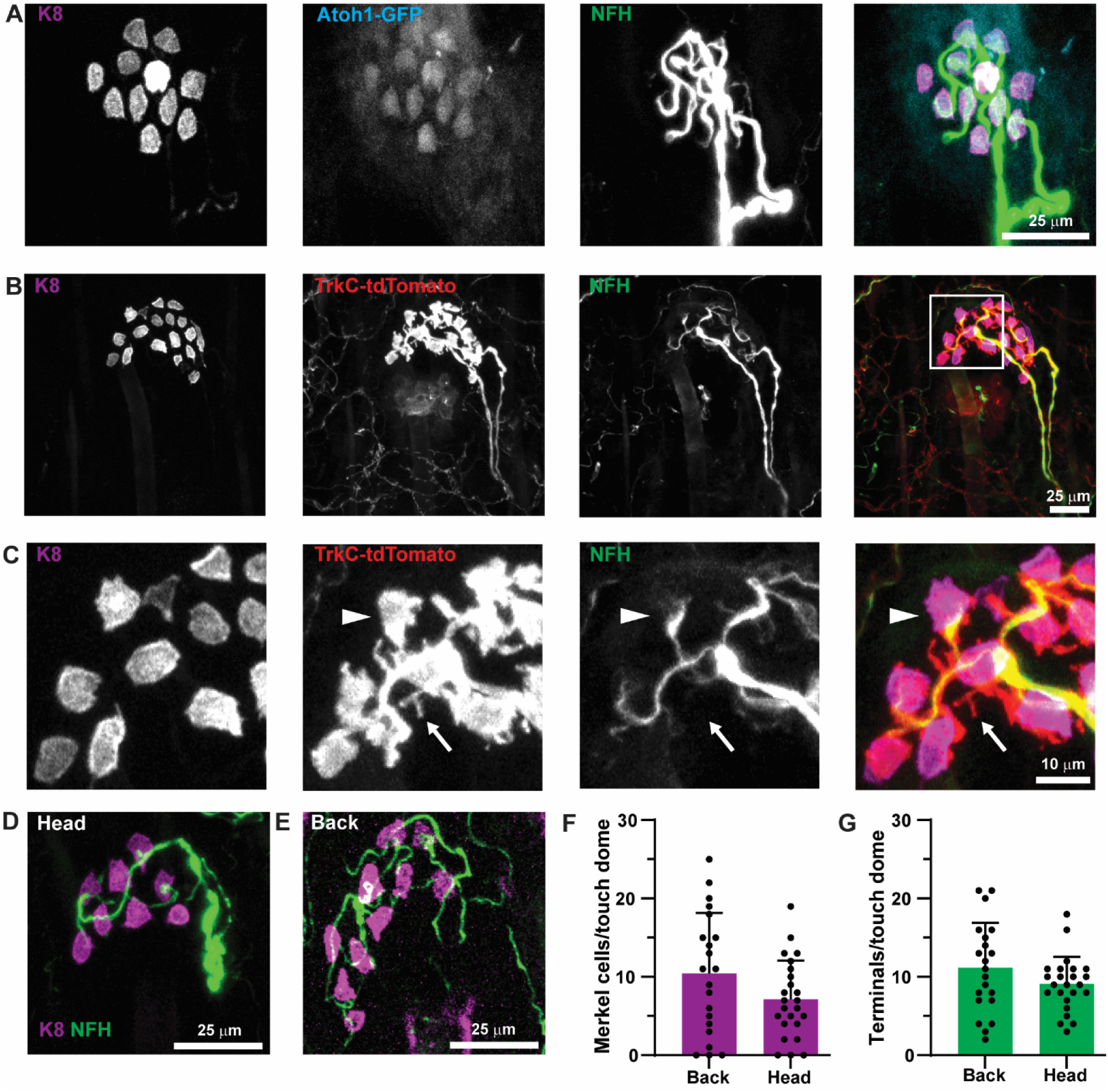
Morphology of mouse Merkel-cell afferents in touch domes. **(A–C)** Representative axial projections of touch domes from *Sarm1+/+*; *Atoh1^A1GFP/A1GFP^*; *TrkC^tdTomato/+^* mouse. Antibodies for (**A**) GFP (cyan, visualized with anti-GFP antibody) or (**B–C**) td-Tomato (red, visualized with anti-DsRed antibody) counterstained with antibodies against K8 (magenta) and NFH (green) to show transgenic label overlap with established markers of Merkel cells and axons. (**C**) Boxed area from (**B**) enlarged to show TrkC-tdTomato endings that make contact with a Merkel cell (arrowhead) and do not contact a Merkel cell (arrow). Note that branches that do not contact a Merkel cell are smaller and do not express NFH at detectable levels. (**D–E**) Representative axial projections of touch domes from skin taken from (**D**) the crown of the head of a *Sarm1^+/+^*; *Atoh1^A1GFP/A1GFP^*; *TrkC^tdTomato/+^* female mouse and (**E**) the lower back of a *C57Bl6/J* female mouse. K8: magenta, NFH: green. (**F–G**) Quantification of the number of Merkel cells (**F**) and terminal branches (**G**) per touch dome. Crown: *N*=24 touch domes, 3 mice (2–12 per mouse); P72–P92. Back: N=21 touch domes, 3 mice (7 per mouse); P66. Welch’s *t* test (two-tail) for Merkel cells (*P*=0.10) or terminal branches (*P*=0.16). Data from back skin were previously published in Marshall et al., (2016; Figure 2B–C).

### Sensory neurons innervating Merkel cells are highly plastic in adult skin

Next, time-lapse *in vivo* imaging of identified Merkel-cell afferents was performed in young adult mice. Anesthetized mice were immobilized using ear bars to stabilize the head for imaging (**Figure 2A**). Three-dimensional Z stacks were acquired with two-photon microscopy to capture high-resolution images of GFP-positive Merkel cells and tdTomato-positive axons. Conventional epifluorescence imaging was used to visualize autofluorescent hair follicles, which served as landmarks to locate and register the same sensory axons across imaging sessions, independent of axonal morphology (**Figure S1**). Axons were imaged through a complete cycle of hair growth, which is approximately one month (**Figure 2B**). Because adjacent hair follicles regenerate synchronously in mice, the hair cycle represents a period of synchronous epithelial-cell turnover. Imaging began when mice were in telogen, the resting phase of the hair cycle, and axons and Merkel-cell clusters were imaged every three days (**Figure 2B**). During the active growth phase anagen, the enlarged hair follicles often obscured the fluorescence of Merkel cells and axons. In such cases, the time point was excluded from quantitative analysis (**Table S1**). A total of 154 images from 20 axons in four mice (4–6 axons per mouse) passed these criteria and were analyzed for plasticity in Merkel cells (**Tables S1–S2**). One axon, which showed dramatic remodeling, was excluded from neuronal plasticity measures as its branch structure could not be registered with confidence across sessions (**Figure S2**; **Table S2**).

**Figure 2.**
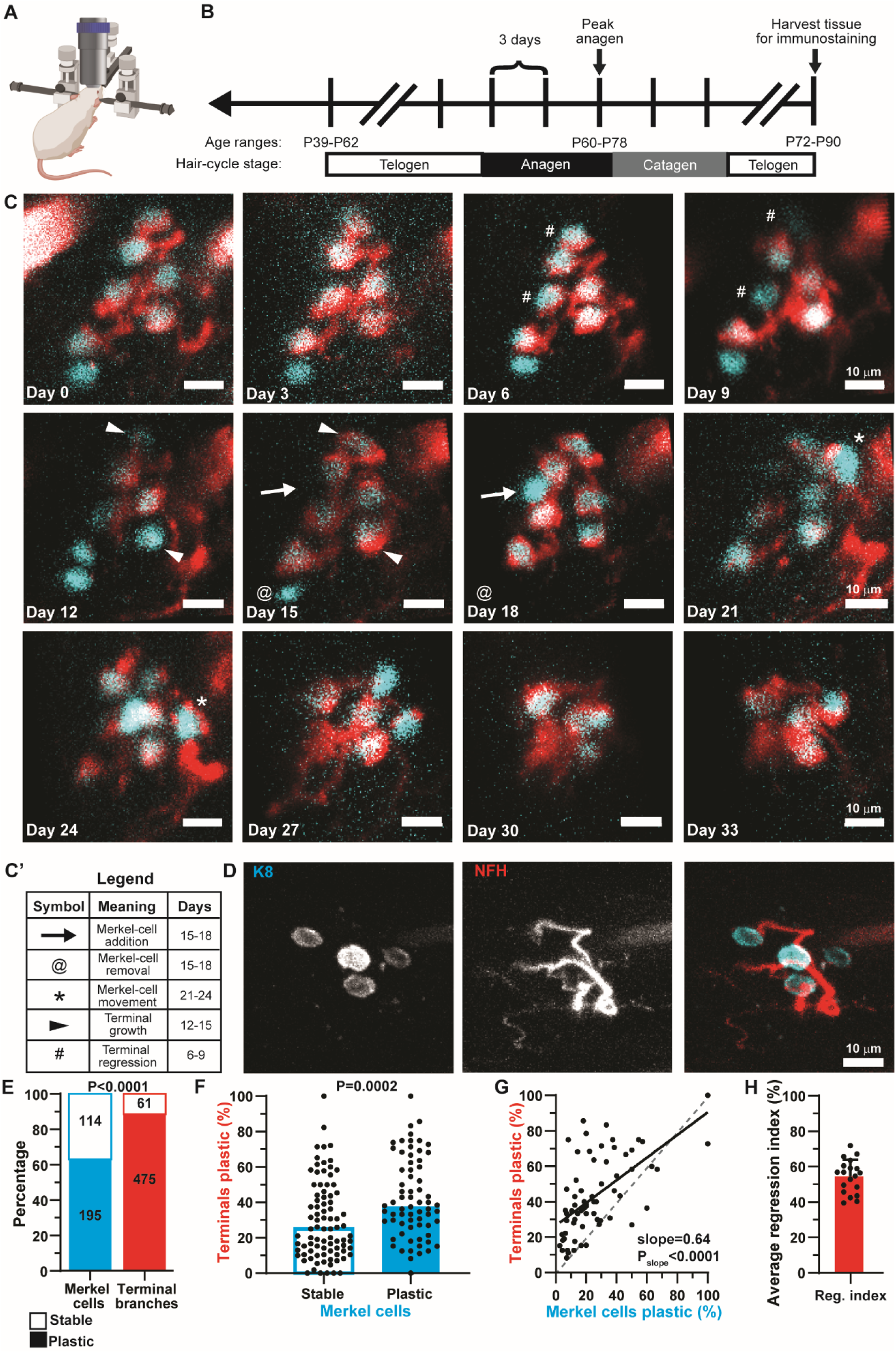
Non-invasive time-lapse imaging reveals that most Merkel cells and axonal terminals show plasticity over 1 month in skin homeostasis. **(A)** Diagram of *in vivo* time-lapse imaging preparation. (**B**) Timeline of experiments, indicating the hair cycle stages throughout imaging and the age ranges at the beginning and end of imaging as well as at peak anagen. (**C**) Representative axial projections of a time series of one *TrkC^tdTomato/+^* arbor (red) and associated *Atoh1^A1GFP/A1GFP^* Merkel cells over the entire duration of *in vivo* time-lapse imaging. Day of image relative to first imaging session indicated in bottom left corner. (**C’**) Legend for symbols used in **C**. Symbols located in the same positions on adjacent days such that the indicated change takes place between those two sessions. (**D**) Axial projection of confocal stack of registered axon from (**C**), labeled with antibodies against K8 (cyan) and NFH (red). (**E**) Comparison of the percentage of Merkel cells and terminals that showed plasticity or were stable throughout the entirety of imaging. *P* value: two-tailed Fisher’s exact test. *N*=309 Merkel cells and 536 terminals from 4 animals. (**F**) Analysis of the percentage of terminals per touch dome per time point showing plasticity when Merkel cells were stable (cyan outline; *N*=86) or showed plasticity (cyan filled; *N*=68). Colored bars indicate median. *P* value: two-tailed Mann-Whitney test. Each dot represents a single time-lapse session from one touch dome (*N*=19 touch domes from 4 mice; 154 total time-lapse sessions). (**G**) Scatter plot of the percentage of Merkel cells and terminal branches showing plasticity in sessions with plasticity in both Merkel cells and terminals. Gray dashed line indicates unity line. Black line is linear regression indicating positive correlation between Merkel-cell and terminal plasticity when both occur. R^2^=0.37, *n*=67 time-lapse sessions. (**H**) Average regression index for each axonal arbor across all time points. *N*=19 arbors; 4 mice. Bar indicates mean±SD.

By comparing the structure of axonal arbors in one session relative to the previous session, we identified several types of plasticity. We tracked the number and relative location of each Atoh1-GFP Merkel cell at each time point and identified Merkel cells that were added, removed, changed locations or remained stable (**Figure 2C–C’**). For TdTomato-positive terminal branches, we noted whether new branches sprouted, and whether existing branches remained stable, were removed, or grew or regressed in length or ending size (**Figure 2C–C’**). After the final imaging session, full-thickness tissue clearing and immunostaining was used to confirm that *in vivo* two-photon imaging sufficed to capture the number and arrangement of Merkel cells and terminal branches observed with our established methods (**Figure 2D**; *N*=12 touch domes from 3 mice).

Overall, both axon terminals and Merkel cells were highly dynamic throughout the imaging period. Plasticity measures did not differ when animals were grouped by sex or age (**Figure S3**). We observed significantly more remodeling events in terminal branches (89%) compared with Merkel cells (63%; **Figure 2E**; *N*=19–20 axons, 4 animals; see also **Figure S3**). This surprising finding suggests that structural plasticity of terminal branches exceeds that of their Merkel-cell targets; therefore, we next tested whether the degree of plasticity in axonal terminals related to that of Merkel cells. By comparing sessions in which Merkel-cell clusters were stable versus plastic, we found that the percentage of plastic terminal branches was increased when Merkel cells were also plastic (**Figure 2F**). Moreover, the degree of plasticity was positively correlated between Merkel cells and terminals in these sessions (**Figure 2G**). Next, we analyzed the balance of axonal growth and regression by calculating a regression index, defined as the percentage of axonal plasticity events that were branch removals or regressions. The average regression index was 55% across arbors (**Figure 2H**; see also **Figure S3**), indicating that branches show almost equal amounts of growth and regression. Together, these data show that axonal arbors are highly dynamic over three-day intervals, and that their amount of structural plasticity correlates in part with changes in Merkel cells.

### Newly specified Merkel cells have short lifetimes

To examine the dynamics of Atoh1-GFP Merkel cells, we quantified instances of addition, removal and relocation in Atoh1-GFP Merkel cells across all sessions (**Figure 3A**). More than half of Merkel cells were removed during the imaging period; however, some appeared to be added or relocated within the cluster, and some Merkel cells underwent multiple plasticity events (e.g., addition followed by movement; **Figure 3B**).

**Figure 3.**
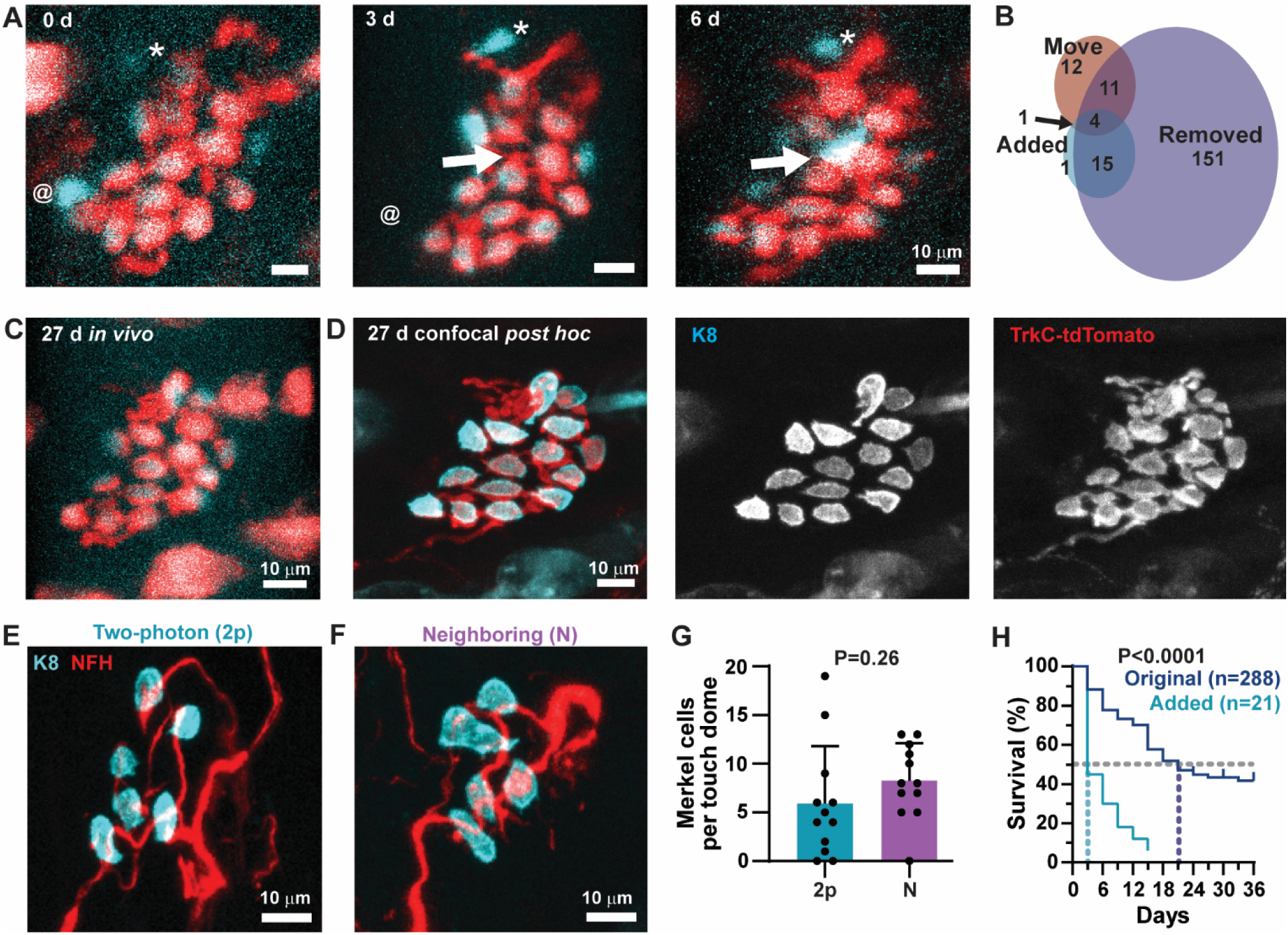
*Atoh1^GFP^* Merkel cells added during adulthood have short lifetimes compared to existing Merkel cells. (**A**) Example axial projections of a touch dome with *Atoh1^GFP^* Merkel cells that were added, removed, and showed presumptive movements. The at symbol denotes a Merkel cell (day 0) that was removed in the next session (day 3). Arrow indicates a Merkel cell that was added on day 6 to the empty spot indicated by the arrow on day 3. Asterisk denotes a Merkel cell moving across sessions (day 0–6). (**B**) Prevalence of each type of plasticity observed in plastic Merkel cells. Areas of overlap indicate Merkel cells that went through multiple types of plasticity. *N*=195 Merkel cells. (**C**) Axial projection of the last in vivo image of the touch dome from (**A**) before the end of time-lapse imaging. (**D**) Axial projection of confocal registration of *in vivo* time-lapse touch dome with antibody staining for K8 (cyan) and DsRed to amplify *TrkC^tdTomato^* (red). (**E**) Example axial projection of confocal-registered touch dome that was part of the time-lapse in vivo dataset with antibodies against K8 (cyan) and NFH (red). (**F**) Example confocal axial projection of neighboring touch dome, which is from the same piece of tissue as **E** but was not imaged during the time-lapse *in vivo* study. (**G**) Comparison of the number of K8-positive Merkel cells in touch domes repeatedly imaged with two-photon imaging (2p) and touch domes that were not subject to repeated irradiation (N). No significant difference in number of Merkel cells per touch dome by Student’s *t* test (*P*=0.26; *N*=12 touch domes in each group from 2–3 animals). Bars indicate mean±SD. (**H**) Survival plot of Merkel cells that were present on day 0 (Original) and Merkel cells that were added at some point during the *in vivo* imaging period (Added). Gray dashed line indicated 50% survival. Colored dashed lines indicate median points for each group. Median survival=3 days for added Merkel cells and 21 days for original Merkel cells. *P* value: Log-rank test to compare survival curves.

The most common type of plasticity observed in Merkel cells was removal (**Figure 3B**); however, we reasoned that what appeared to be removal could be either a change in the level of fluorescent reporter expression or could be removals induced by photodamage from repeated two-photon irradiation. We confirmed that the loss of Atoh1-GFP signal did not simply reflect downregulation of *Atoh1* expression by registering Atoh1-GFP-positive Merkel cells in the last session of *in vivo* imaging (**Figure 3C**) with *post hoc* immunofluorescence staining for the mature Merkel-cell marker K8 (**Figure 3D**). We observed almost complete overlap between these Merkel-cell markers (71/72 K8-positive Merkel cells also expressed Atoh1-GFP), indicating that loss of Atoh1-GFP signal reflects the disappearance of a Merkel cell. We next tested whether photodamage accounts for the loss of Merkel cells observed throughout time-lapse imaging studies. The number of K8-positive Merkel cells in touch domes that were repeatedly imaged *in vivo* with a two-photon laser (**Figure 3E**) was compared with the number of Merkel cells in neighboring touch domes that were not subjected to two-photon irradiation (**Figure 3F**). The number of Merkel cells per touch dome was not significantly different between these two groups (**Figure 3G**), suggesting that the decline in Merkel-cell numbers is not due to photodamage from two-photon imaging. Thus, our results indicate that Merkel-cell numbers decline substantially over the course of a month in adult mice.

In addition to removals, 7% of all Merkel cells appeared to be added and 9% relocated within the cluster. Given that Merkel cells are derived from non-migrating keratinocyte precursors (Doucet et al., 2013; Xiao et al., 2015), we were surprised to see that 14% of plastic Merkel cells appeared to change locations during imaging. It is possible that a Merkel cell was removed and a new Merkel cell was added in a nearby location; however, scoring a single Merkel cell as relocated, rather than the removal of one cell and the addition of a new cell, is a more conservative measure of plasticity, since Merkel cells are reported to be long-lived compared to keratinocytes (Doucet et al., 2013; Wright et al., 2017). Merkel cells were scored as relocated when their cell shape was the same as a cell in the previous session and a line could be drawn between the previous location and the new location without crossing other Merkel cells. We found that almost all movements were coincident with or after other Merkel cells were removed from the arbor (n=32/35 movements). Moreover, Merkel cells relocated toward either the center of the cluster or a position previously occupied by another Merkel cell. Together, these results suggest that removal events can trigger the rearrangement of remaining Merkel cells within the axonal arbor.

Finally, we noted that 10% of Merkel cells went through multiple plasticity events, for example added Merkel cells were almost always removed before the end of the imaging period (**Figure 3B**). We compared the survival curves of added Merkel cells with those of original Merkel cells that were present during session 0 and found that most added Merkel cells turned over within one imaging session whereas most original Merkel cells persisted for longer than 18 days (**Figure 3H**). These data confirm a previous report that adult-born Merkel cells have short lifetimes when compared with resident Merkel cells in ventral skin (Wright et al., 2017). These results suggest that newly specified Merkel cells are not stably incorporated into axonal arbors over a one-month time frame.

### Sensory axon terminals targeting Merkel cells have two distinct morphologies

Given that we observed two populations of Merkel cells with distinct lifetimes, we wondered whether the contacts between Merkel cells and axonal terminals were also heterogeneous. The canonical Merkel disc is a shallow cup that encloses only the lower surface of the Merkel cell (Iggo & Muir, 1969), and terminals with this morphology were commonly observed *in vivo* (**Figure 4A–B**). To our surprise, we also noted endings that lacked the large cup and instead terminated in a modest swelling (**Figure 4A–B**), which has not been previously described in axon terminals contacting Merkel cells. To test whether these structures represent two distinct populations, we measured the length and width of every ending at each time point that a Merkel cell was contacted and approximated the ending area as an ellipse (**Figure 4C–D**). In a histogram of the maximum ellipse area of each ending, we noted a distribution that was best fit by bimodal gaussian (**Figure 4E–F**; R^2^=0.80, bimodal; R^2^=0.76, unimodal distribution), suggesting that the smaller endings are a distinct population, rather than the trailing edge of a continuum. We refer to these previously undescribed endings as *boutons*, which are typical of many axons throughout the nervous system. Inspired by calyx endings of vertebrate hair cells, we term the Merkel cell’s canonical terminal structure *kylix* (plural: *kylikes*) after the Greek wine cup whose shape is a shallow, stemmed bowl (**Figure 4G**). To estimate a size threshold to distinguish boutons and kylikes, the component gaussians of the bimodal fit were plotted and the point at which the two component curves crossed was defined as the threshold between the two structures (**Figure 4F**; gray dashed line; threshold=16.3 µm^2^). Together, these data suggest that Merkel cells are contacted by axonal branches with two distinct sizes.

**Figure 4.**
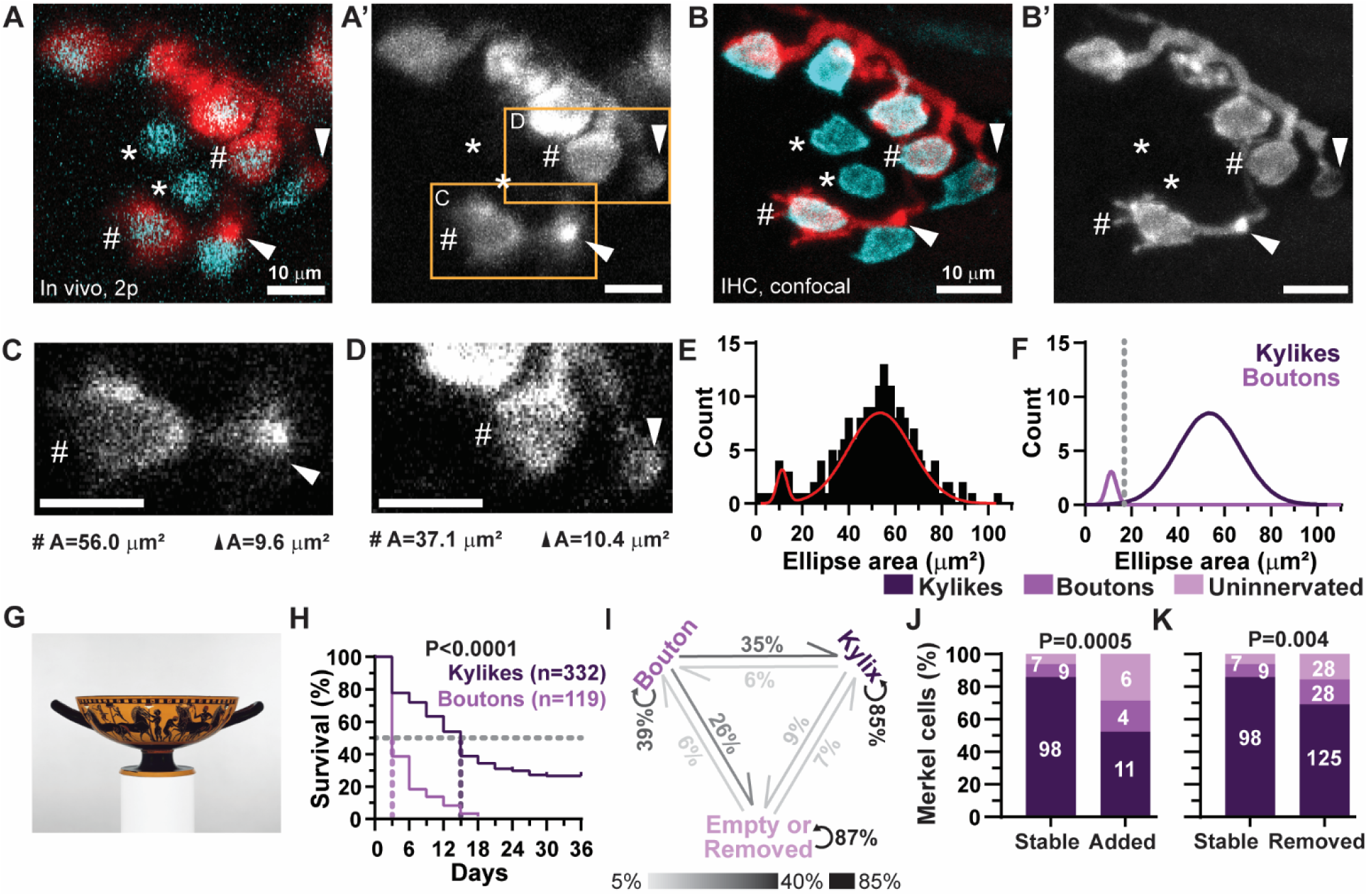
Merkel cells are contacted by two morphologically distinct types of afferent terminals. (**A–A’**) In vivo image of touch dome with (**A**) and without (**A’**) Merkel-cell overlay displaying all three contact types. Symbols indicate examples of each ending type: asterisks, uninnervated; hashtags, kylikes; arrowheads, boutons. (**B–B’**) Confocal image of same touch dome as **A**, labeled with antibodies against dsRed (TrkC^tdTomato^, red) with (**B**) and without (**B’**) K8 (cyan) overlay. Symbols as in **A**. (**C–D**) Single z planes of zoomed in regions from **A’**, used to measure length and width of individual endings. Areas for each ending indicated below image. (**E**) Histogram of the maximum ellipse area for each identified terminal, so that each terminal is represented once. Red line: best fit bimodal gaussian. (**F**) Gaussian components underlying best fit bimodal gaussian in **E**. Threshold (gray dashed line) defined as the crossing point between the two component gaussians. (**G**) Ancient Greek kylix. Note the shallow cup and thin stem of the kylix, which closely resembles the previously described Merkel-cell contact. Image from the Metropolitan Museum of Art, reproduced under Creative Commons license CC0 1.0. (**H**) Survival plot comparing boutons and kylikes. *P* value: Log-rank test. *N*=339 kylikes and 120 boutons. (**I**) Schematic indicating the probability of morphological changes in terminal branches that contact Merkel cells. *N*=1,120 kylikes, 153 boutons, and 168 empty terminals. The most probable event for each ending type was for the morphology to remain the same (kylikes: 85%, boutons: 39%, empty/removed endings: 87%). (**J**) Comparison of stable and added Merkel cells with the first known contact of each type. *P* value: two-tailed Fisher’s exact test. (**K**) Comparison of the percentage of stable and removed Merkel cells with the last known contact of each type. *P* value: two-tailed Fisher’s exact test.

As kylikes are more abundant than boutons, we hypothesized that the former are longer lived structures. To test this hypothesis, we calculated the persistence length of each ending that contacted Merkel cells. Kylikes showed a fivefold longer median survival time than boutons, demonstrating that they are indeed a more stable structure (**Figure 4H**).

We next asked whether the kylix and the bouton are independent structures, or whether individual terminal branches could transition between these morphologies. The most probable event for each type of terminal branch was to maintain its morphological subtype between successive imaging sessions; however, each type of terminal branch had some changes in ending morphology. Kylikes were rarely observed changing morphologies; only 6% of kylikes regressed to boutons and 9% became either empty branches (without a Merkel-cell contact) or were completely removed along with their terminal branch. By contrast, boutons changed morphologies in >60% of observations: 35% of transitions were growth into kylikes, and 26% of transitions were to empty branches that lacked Merkel-cell contacts or were completely removed (**Figure 4I**). Thus, we conclude that boutons are an intermediate structure observed during both terminal branch growth and regression.

Since axon terminals differed in both size and stability, we wondered whether the contact types differed between stable and plastic Merkel cells. We compared the last known contacts associated with stable Merkel cells with those of added (**Figure 4J**) and removed (**Figure 4K**) Merkel cells. Compared to stable Merkel cells, both added and removed Merkel cells were more likely to be either uninnervated or contacted by a bouton (**Figure 4J–K**). Together, these data show that stable Merkel cells are highly likely to be innervated by long-lived kylikes, whereas plastic Merkel cells may be either uninnervated, contacted by short-lived boutons, or innervated by kylikes.

### Axonal plasticity is stochastic but Merkel-cell remodeling is synchronized to hair growth

We next analyzed the temporal dynamics of structural plasticity in Merkel cell-neurite complexes. For each imaging session, we calculated a plasticity index, which is the percentage of Merkel cells in each arbor that showed a structural change from the previous session (**Figure 5A**). The plasticity index of terminal branches was likewise calculated on a per-arbor basis (**Figure 5B**).

**Figure 5.**
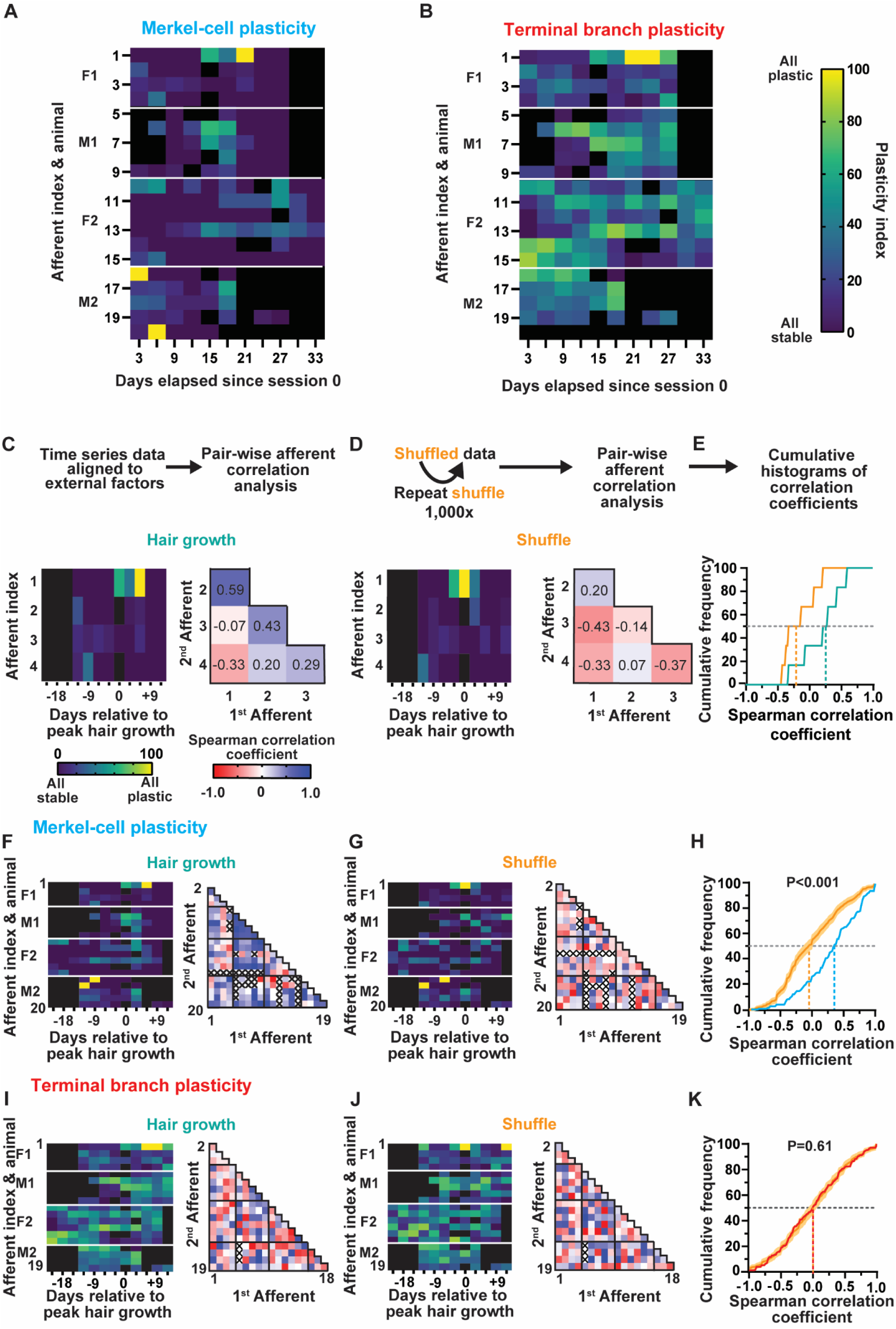
The plasticity of Merkel cells but not axon terminals is synchronized across arbors by the hair cycle. (**A–B**) Plots indicating the Merkel-cell (**A**) and terminal branch (**B**) plasticity indices for each afferent on each session. Color scale in **B** applies to both **A** and **B**. White lines mark afferents belonging to each animal. Black squares indicate sessions with missing or unanalyzable data. (**C–E**) Schematic of analysis pipeline to test similarities in temporal patterns of plasticity across afferents. (**C**) Time series data from animal F1 are aligned to external factors, e.g. hair growth. Then, a correlation matrix is generated to calculate a correlation coefficient for each pair of afferents. (**D**) To test whether correlations are higher than chance, data are shuffled 1,000 times and a correlation matrix is generated for each set of shuffled data. (**E**) Correlation coefficients from experimental (**C**) and shuffled (**D**) datasets are compared using cumulative histograms. Gray dashed line indicates 50%, colored dashed lines indicate median correlation coefficients for each group. (**F–K**) Analysis of Merkel-cell plasticity index (**F–H**) and terminal branch plasticity index (**I–K**) for every afferent used in analysis following the same organization as **C–E**. (**H,K**) Cumulative histograms of correlation coefficients from experimental data and mean±SD cumulative distribution of shuffled permutations. *P* values: permutation test.

Both Merkel cells and axonal branches showed several temporal patterns of plasticity. For example, 6/20 of Merkel-cell clusters showed prolonged periods of plasticity (>10% plasticity for ≥3 consecutive sessions). In five of these arbors, Merkel-cell clusters showed dramatic remodeling in which ≥50% of cells changed between two consecutive sessions (e.g., afferents 1 and 11). Other clusters showed sporadic plasticity interspersed with sessions of stability. Additionally, every Merkel-cell cluster showed periods of stability that ranged from 1–9 sessions (**Figure 5A**). Conversely, axonal branches were more likely to remodel continuously than were Merkel-cell clusters (**Figure 5B**). Together, these data suggest that remodeling is continuous in axonal arbors but can be either sporadic or ongoing in Merkel-cell clusters.

Next, we wondered whether periods of remodeling and stability were synchronized across arbors by biological or extrinsic factors. We identified two candidate factors that could govern remodeling of touch receptors in the skin: the hair-growth cycle and two-photon irradiation. During hair growth, skin undergoes synchronous changes in cellular proliferation and innervation density (Muller-Rover et al., 2001; Botchkarev et al., 1997; Peters et al., 2001), thus the hair cycle could provide signals that modulate plasticity in Merkel cell-neurite complexes. Additionally, we reasoned that two-photon irradiation could damage the skin and induce plasticity in response. To test whether either hair growth (peak anagen, **Figure 5**) or amount of two-photon irradiation (two-photon session number; **Figure S4**) synchronized plasticity across arbors, we aligned temporal trajectories of plasticity parameters (Merkel-cell plasticity index, terminal branch plasticity index) to each factor. Then, we generated a correlation matrix, which calculates Spearman correlation coefficients for all possible pairs of arbors (**Figure 5C**). To test whether these correlation coefficients were different from what would occur by chance, we shuffled the order of time-series data 1,000 times and then performed correlation analysis (**Figure 5D**). Shuffled and experimental correlation coefficients were compared with cumulative histograms (**Figure 5E**); *P* values were calculated with the permutation test, which asks whether the experimental median correlation coefficient falls within the distribution of medians from shuffled datasets. We applied this analysis method to plasticity indices for Merkel cells (**Figure 5F–H**) and axon terminals (**Figure 5I–K**). When aligned by two-photon session number, neither Merkel-cell nor axonal plasticity was correlated across arbors (**Figure S4**). By contrast, when data were aligned to peak hair growth, Merkel-cell plasticity was significantly correlated across arbors (**Figure 5H**), whereas the plasticity of axon terminals was asynchronous (**Figure 5K**). These surprising results indicate that Merkel-cell plasticity, but not axonal remodeling, is synchronized to the rapid epithelial regeneration that occurs during hair growth.

### Axonal branches are stabilized by Merkel-cell contacts

Since plasticity in Merkel cells and axon terminals are only loosely correlated, we wondered how connections with Merkel cells impact branch plasticity. To address this question, we tested whether plasticity in branches that contacted Merkel cells (“occupied”) differed from plasticity in branches that never contacted a Merkel cell during the imaging period (“empty”; **Figure 6A–B**). All empty branches were plastic whereas 20% of occupied branches were stable throughout the imaging period (**Figure 6C**). Additionally, empty branches showed a significantly shorter lifetime than branches that contacted Merkel cells (**Figure 6D**). Since Merkel cells moderately stabilize branches, we next asked what happens to branches after Merkel cells are removed. Surprisingly, only 50% of branches were removed on the same day as the Merkel cell. An almost equal number of branches (47%) were either removed after the Merkel cell or persisted until the end of imaging (**Figure 6E**). Together, these analyses show that Merkel-cell contacts bias branches toward stability, but other stability-promoting mechanisms must also exist.

**Figure 6.**
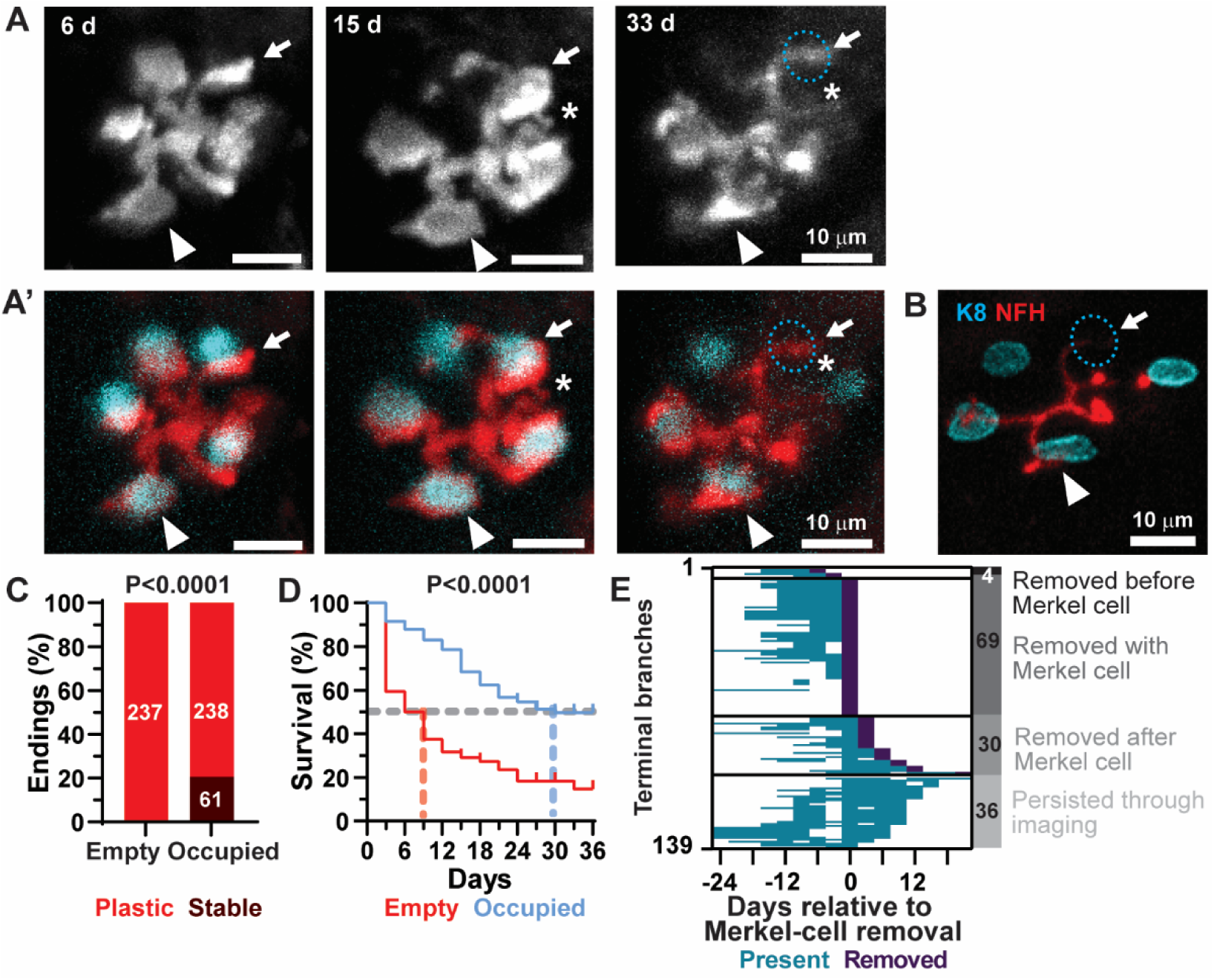
*TrkC^tdTomato^* afferent terminals are more stable if they contact Merkel cells. (**A–A’**) Axial projections of *TrkC^tdTomato^* terminals of one identified afferent over time without (**A**) and with (**A’**) Merkel-cell overlay. Days elapsed since day 0 indicated in top left corner. (**B**) Axial projection of confocal registration with antibodies against K8 (cyan) and NFH (red). Arrowhead, stable occupied branch; arrow, occupied branch that persists after Merkel-cell removal; blue dashed circle, location of removed Merkel cell. Symbols located in the same position on adjacent days indicate change occurring between those two sessions. All changes were noted in three-dimensional stacks. (**C**) Comparison of the percentage of Merkel-cell contacting (occupied) and empty terminals that were stable for the duration of imaging or showed plasticity. *P* value: two-tailed Fisher’s exact test. (**D**) Survival curve comparing empty and occupied terminal neurites. Gray dashed line indicates 50% survival. Colored dashed lines indicate median values for each group (9 d for empty terminals and 30 d for occupied terminals). *P* value: log-rank test. (**E**) Trajectories of individual branches associated with removed Merkel cells aligned to the day of Merkel-cell removal. Bar on right indicates how many branches were either not removed (persisted through imaging) or removed before, with, or after Merkel-cell removal.

We next asked whether Merkel-cell removal impacts branch plasticity across an entire arbor (**Figure 7A–C**). The number of sprouting branches was compared one session after instances where Merkel cells were and were not removed. Significantly more branches were added in the session after a Merkel cell was removed than after sessions that lacked Merkel-cell removal (**Figure 7D**). Additionally, instead of terminating in the Merkel-cell layer as is typical of Merkel-cell afferents (**Figure 7E**), some sprouting branches projected to the superficial levels of the epidermis (**Figure 7F**), akin to the overshoot process seen during development of the Merkel cell-neurite complex (Jenkins et al., 2019). In total, all animals had an arbor with overshoot at some point during imaging, with half of all arbors having at least one overshooting branch (**Figure 7G**). These overshooting branches most frequently occurred either in the same session or in sessions after Merkel-cell removal (**Figure 7H**). Thus, Merkel-cell removal promotes axonal branching and abnormal patterning.

**Figure 7.**
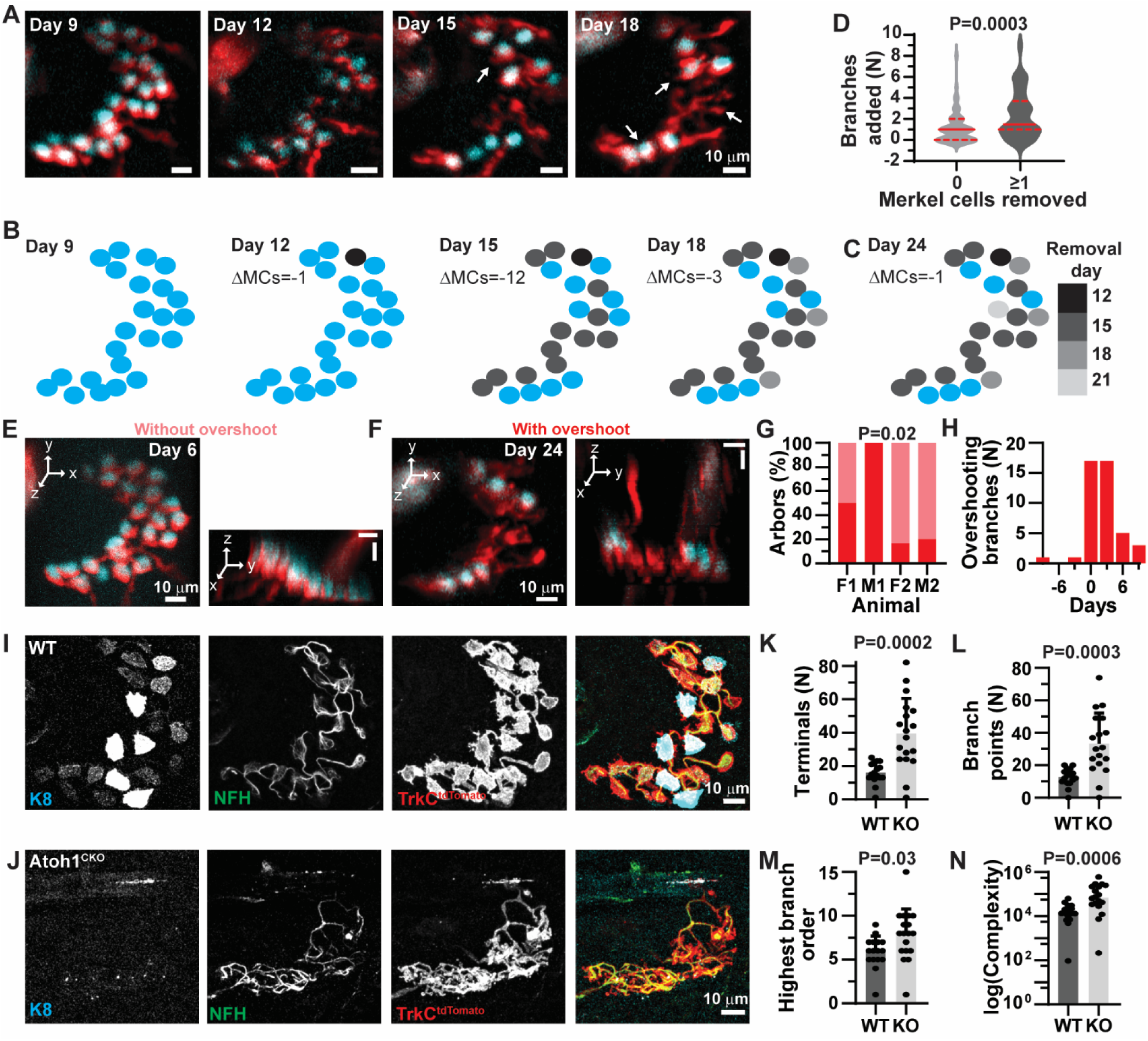
Merkel cells limit branch number and promote branch structural maturation. (**A**) Axial projections of touch dome where there was an increase in the number of branches added after Merkel-cell removal. (**B–C**) Schematic of Merkel cells from Day 9 image in **A**, color coded to indicate which Merkel cells are removed on each day. Color code in C applies to B and C. (**D**) Comparison of the number of branches added in the session after Merkel cells were (≥1) or were not (0) removed. *P* value: Mann-Whitney rank test. (**E–F**) Example z projections (left) and rotated three-dimensional projections (right) from the same arbor, in a session without overshoot (**E**) and with overshoot (**F**). Day of imaging is indicated in upper right corner. Scale bars indicate 10 microns. Stack from **F** is thicker than **E**, thus rotated image is thicker. (**G**) Comparison of the percentage of afferents from each animal displaying overshoot. *P* value is result from Fisher’s exact test (*N*=4–6 touch domes per animal). (**H**) Analysis of timing of branch overshoot relative to the nearest event of Merkel-cell removal. Positive values indicate days after Merkel cell is removed; negative values indicate days before Merkel-cell removal. (**I–J**) Axial projections of touch domes from WT (*K14^+/+^; Atoh1^fl/+^; TrkC^tdTomato/+^*) and Atoh1^CKO^ (*K14^Cre/+;^ Atoh1^fl/lacz^; TrkC^tdTomato/+^*) 8 w animals stained for K8 (cyan), NFH (green) and DsRed to amplify TrkC-tdTomato (red). (**K–N**) Comparison across genotypes of branching parameters: number of terminal branches (**K**), number of branch points (**L**), highest branch order (**M**), and complexity index (**N**). *P* values: Welch’s *t* test (**K–L**), Student’s *t* test (**M**), or Mann-Whitney rank test (**N**). *N*=16–18 touch domes from 2 mice; 7–9 touch domes per mouse. Bars indicate mean±SD (**K‒M**) or median (**N**).

Two models could explain why branches sprout after Merkel cells are removed: either the physical process of Merkel-cell removal stimulates branching or a stable Merkel-cell contact suppresses additional branching. To distinguish between these two models, we used quantitative immunohistochemistry in full thickness skin to analyze the branched structures of arbors from animals that lacked Merkel cells (*K14^Cre/+^; Atoh1^fl/lacz^; TrkC^tdTomato/+^*) and littermate controls (*K14^+/+^; Atoh1^fl/+^; TrkC^tdTomato/+^*; **Figure 7I–J**). Arbors from mice that lacked Merkel cells showed increased terminal branches and branch points, highest branch order, and complexity (**Figure 7K–N**). Additionally, branches in Merkel-cell knockout animals lacked kylikes (**Supplemental Figure 6**). Collectively, these results demonstrate that the presence of Merkel cells, rather than the act of Merkel-cell removal, suppresses branching and promotes structural maturation to the kylix.

## Discussion

To maintain a functional barrier our sensory epithelia must continually renew; however, epithelia are also densely innervated to serve as sensory interfaces, which raises the question of whether sensory axons remodel alongside renewing epithelia. Although this is a fundamental question in sensory neurobiology, little is known about the fidelity with which sensory neurons maintain patterning during epithelial turnover. We used a novel, noninvasive imaging preparation to directly show that both Merkel cells and axon terminals go through common plasticity events during normal tissue homeostasis. Surprisingly, axon terminals frequently remodeled without accompanying changes in their target Merkel cells, suggesting a role for intrinsic neural programs in structural plasticity. On the other hand, Merkel cells stabilize individual branches, suppress exuberant branching and promote maturation of terminal morphology, highlighting the contribution of epithelial-neural crosstalk. Thus, we propose a model in which intrinsic neural mechanisms drive stochastic sprouting and regression of terminal branches. Upon contacting Merkel cells, the probability of branch growth/regression decreases as they mature to the highly stable kylix morphology. The presence of stable Merkel cell-kylix complexes in turn suppresses sprouting of nearby branches (**Figure 8**).

**Figure 8.**
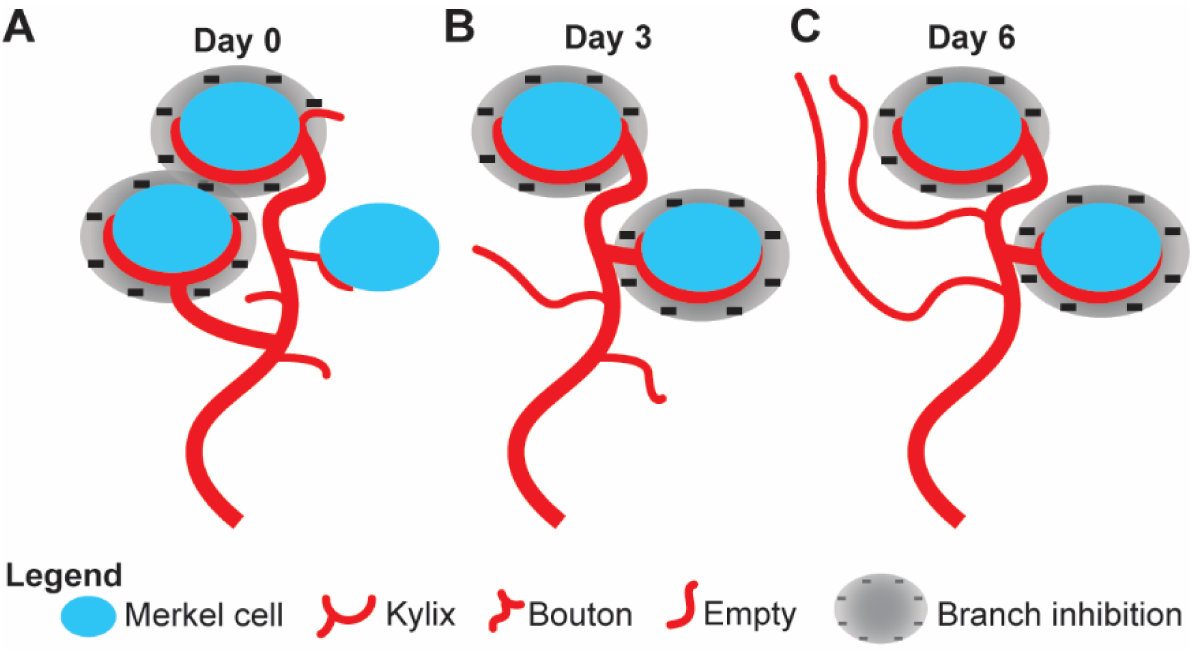
A model of how sensory axons respond to Merkel-cell plasticity. (**A**) Three Merkel cells are contacted by either a kylix or a bouton. Merkel cell-kylix connections inhibit surrounding branches. (**B**) One Merkel cell is removed. The bouton grows into a kylix and inhibits surrounding branches. (**C**) Branches are added and lengths increase such that branches overshoot the Merkel-cell level after a Merkel cell-kylix complex is removed. Legend indicated at the bottom.

### *In vivo* imaging captures structural plasticity of sensory axons during tissue homeostasis, development, injury and disease

Our study of Merkel-cell afferents in adult skin homeostasis provides an in-depth view of the types and prevalence of plasticity events in a homogenous population of mechanosensitive afferents that synapse onto epithelial cells. These studies are the first to our knowledge to simultaneously track both presynaptic epithelial cells and postsynaptic axons, providing a unique perspective on how turnover of epithelial cells affects the sensory afferents targeting those same cells. It is worth noting that, because Merkel-cell afferents in touch domes have punctate receptive fields, our *in vivo* imaging preparation allowed us to track structural remodeling of an afferent’s entire mechanosensory receptive field, ∼10^4^ µm^2^. In the axial dimension, we typically imaged from the stratum corneum to the papillary dermis (∼30 µm below the skin’s surface). Thus, although Merkel-cell afferents are Aβ fibers with thickly myelinated axons, our quantitative analysis focused primarily on the axons’ unmyelinated branches that project into the epithelial compartment and are the site of mechanotransduction.

Although the plasticity of peripheral sensory neurons is understudied, remodeling of central axons and dendrites has been studied in the context of learning and memory. The morphology of dendritic spines is an anatomical correlate of synaptic strength, with spines changing from thin filipodia to enlarged mushrooms as they stabilize. *In vivo* longitudinal imaging in adult mice has revealed that, although the gross morphology of axons and dendrites is stable over several weeks, both dendritic spines and a subset of axonal boutons show structural plasticity (Trachtenberg et al., 2002; De Paola et al., 2006; Grillo et al., 2013). Terminal branches imaged in our study had a higher rate of turnover than that previously observed in cortical axons: 70% of kylikes, the most stable ending type in our study, turned over within 21 d, whereas only 40% of terminal boutons turned over during the same time period in a study of axonal dynamics in somatosensory cortex of adult mice (Grillo et al., 2013). The increased remodeling of axons imaged during our study could be required to maintain healthy axonal endings in a tissue that is both renewing and located superficially, which subjects axon terminals to physical stresses and strains.

The neuromuscular junction is another peripheral site where axons interact with non-neuronal target cells. Similar to axons in the central nervous system, motor axons go through a process of outgrowth, competition and pruning during development (Balice-Gordon & Lichtman, 1993; Walsh & Lichtman, 2003). By contrast with our findings, the morphology of mature neuromuscular junctions remains stable throughout adulthood (Lichtman et al., 1987; Balice-Gordon & Lichtman, 1990). Thus, although the neuromuscular junction shares some similarities with the Merkel cell-neurite complex, these are fundamentally different end organs with distinct dynamics.

Our findings of axonal plasticity during skin homeostasis complement recent studies that used longitudinal *in vivo* imaging of vertebrate sensory axons in other contexts. For example, a recent report showed that the unmyelinated branches of a heterogeneous population of nociceptors grow and retract on the order of 5 µm per hour *in vivo* (Takahashi et al., 2019). These changes are on a similar scale to those we observed in terminal branches of over three days, supporting the notion that ongoing remodeling is stochastic. Interestingly, the same report found that axonal pruning is reduced in a mouse model of atopic dermatitis. Together with our findings, these results suggest that hyperinnervation observed in inflammatory skin conditions reflects the process of homeostatic neuronal plasticity coupled with attenuated branch pruning.

Numerous studies have demonstrated the regenerative capacity of sensory axons after injury. Most studies have focused on axonal growth *in vitro*, or at defined time points across specimens *in vivo* (Mahar & Cavalli, 2018; Waller, 1850; Lunn et al., 1989; Osterloh et al., 2012); however, a handful of *in vivo* studies have tracked the same nerves over time. In studies utilizing laser ablation and time-lapse imaging of single axonal branches in the skin, peripheral axons degenerated within 6 days of lesioning and showed regrowth 10 days after lesioning (Yuryev & Khiroug, 2012; Yuryev et al., 2014). Another commonly used model to study axonal injuries is the spared nerve injury model of neuropathic pain. This technique was recently applied in combination with *in vivo* imaging to demonstrate that nociceptive axons, but not touch receptors, sprout collaterals over millimeters to re-innervate skin after injury (Gangadharan et al., 2022). For these experiments, heterogeneous groups of touch receptors and nociceptors were imaged and overall fiber length was computed. In sham operated controls, no difference in this metric was observed. This finding is not surprising, as overall length is dominated by axonal bundles traversing millimeters, which overshadows any changes in axon terminals at the site of initial sensory encoding. It has been long known that axons in the central nervous system do not regrow after injury; however, *in vivo* time lapse imaging has revealed that central-projecting branches of sensory axons do display some regeneration after injury. As opposed to peripheral axonal branches, spinal cord projecting branches show modest regeneration, with fewer branches and slower regeneration when compared with peripheral branches. Moreover, central branches showed pathfinding errors and failed to regrow through the injury site to the original target (Kerschensteiner et al., 2005).

Non-invasive imaging of zebrafish embryos has also identified mechanisms of remodeling during development of sensory systems. Lateral-line neuromasts are epithelial structures comprising mechanically sensitive hair cells innervated by sensory afferents (Dow et al., 2018; Dow et al., 2015; Erzberger et al., 2020). Within a neuromast, a given afferent precisely targets all hair cells with the same directional selectivity in a Notch-dependent manner (Dow et al., 2015, Dow et al., 2018). Interestingly, hair cells transiently grow projections that assist in guiding afferents to mature synaptic sites (Dow et al., 2015). Similarly, some Merkel cells show a dendritic-like morphology with protrusions (Kim & Holbrook, 1995). Future studies with finer temporal precision are needed to test whether these protrusions aid in recruiting afferent terminals to form kylikes. Live-cell imaging of zebrafish embryos has also been used to study development of peripheral axon arbors in Rohon Beard cells, a population of gentle-touch neurons that transiently exist during development and are responsible for initiating escape responses (Clarke et al., 1984; Haynes et al., 2022). Recent work has shown that peripheral arborization of Rohon Beard cells depends on kinesin light chain 4 (Haynes et al., 2022), which regulates binding between cellular cargo and kinesin motors (Morfini et al., 2016). In the absence of KLC4, new branches initiate but do not properly separate from sister branches, which ultimately results in abnormal axon fasciculation (Haynes et al., 2022). This disorganized phenotype is reminiscent of the hyperbranched and tangled afferents from mice lacking Merkel cells. An intriguing possibility is that these morphological deficits are due to impaired transport of cellular cargos.

### Merkel cells are dynamic during adult skin homeostasis

Previous studies have documented turnover of Merkel cells in adult skin. Genetic lineage tracing suggests that mouse Merkel cells from dorsal skin have lifetimes of approximately eight weeks (Doucet et al., 2013). A more recent *in vivo* study that tracked individual Merkel cells in ventral skin indicates that more than half of these Merkel cells persisted for 13 weeks in the absence of barrier disruption (Wright et al., 2017). In both our study and this previous study, newly added Merkel cells had dramatically shorter lifetimes compared with Merkel cells observed in the first imaging session. Moreover both studies found that new Merkel cells were less likely to become innervated by kylikes, which were observed in living tissue here versus *post hoc* analysis of NFH staining in fixed tissue (Wright et al., 2017). Interestingly, the previous study found that the rate of Merkel-cell production in adult ventral skin offsets Merkel-cell losses to maintain the size of Merkel-cell clusters. Whereas we observed a similarly low rate of Merkel-cell production in the crown of the head, cluster sizes decreased over time due to a higher rate of Merkel-cell removal. In agreement with our findings, other groups have documented age-related loss of Merkel cells in mice and humans (Feng et al., 2018; Moayedi et al., 2018; Garcia-Piqueras et al., 2019). Overall, the results of these two *in vivo* studies are in excellent agreement, and minor discrepancies might reflect methodological differences, strain-specific effects or biological differences between body sites, since Merkel-cell clusters are larger in ventral skin (∼30 Merkel cells per cluster) than those on the crown of the head (14±6 Merkel cells per cluster at the initial imaging session).

### Axonal terminals form two distinct types of morphological contacts with Merkel cells

We found two morphologically distinct axon terminals, kylikes and boutons, that make contacts with Merkel cells. The morphology of the kylix has long been described as the canonical Merkel-cell contact (Iggo & Muir, 1969). We noted that this structure, with a large cup making contact with presynaptic cells, resembled the calyceal endings of vestibular type 1 hair cells and the Calyx of Held (Held, 1893). Whereas calyces engulf the entire target cell, axons targeting Merkel cells formed shallower cups that only touch the lower surface of the cell. Thus, we termed the Merkel-cell contact a kylix, after the drinking vessel with a shallow cup and thin stem.

The smaller axonal ending, referred to here as a bouton, has only been described in frogs. These endings lack morphological specializations of synapses, contained dense core vesicles, and showed sparse mitochondria (Bani, 2009). By contrast, kylikes are rich in mitochondria and have clear core vesicles (Iggo & Muir, 1969; Chen et al., 1973; Mihara et al., 1979). The previous report hypothesized that boutons are efferent endings (Bani, 2009); however, we found that both kylikes and boutons are formed from TrkC-positive sensory axons. Thus, boutons are afferent endings along with kylikes. Additionally, we show that boutons are capable of both growing into kylikes and regressing to a branch without a synaptic contact or complete branch removal, thus we propose that boutons are an intermediate structure for both branch growth and regression. Future studies will investigate whether there are differences in the expression of mechanically activated ion channels and synaptic molecules in kylikes and boutons.

### Merkel-cell plasticity, but not axon plasticity, is synchronized during epithelial turnover

We were surprised to find that Merkel cell remodeling, but not axonal remodeling, was synchronized during active hair growth, which is a period of high epithelial cell turnover. Previous histochemical studies from our group and others have shown that Merkel cells and innervation density fluctuate during hair growth cycles (Marshall et al., 2016; Moll et al., 1996; Nakafusa et al., 2006). By immunolabeling and reconstructing Merkel-cell afferents in full-thickness skin specimens, we observed that, during hair growth, the number of K8+ Merkel cells per axonal arbor decreased, and NFH+ axonal endings simplified (Marshall et al., 2016). This previous study used antibody labeling to examine sensory axons and Merkel cells via the intermediate filament proteins NFH and K8, respectively. Here, live imaging studies utilized the transgenic reporter TrkC-tdTomato, which labels the cytosol rather than the intermediate filament cytoskeleton. Surprisingly, more axonal branches were labeled by TrkC-tdTomato than NFH antibodies. Branches that were TrkC-tdTomato+/NFH- tended to form either boutons or small terminals that did not contact a Merkel cell. Thus, this population was not detected by our previous methods and might contribute to differences in Merkel-cell and axon plasticity observed in our studies. Additionally, we previously showed that the density of epidermis-innervating NFH+ axons increased during active hair growth (Marshall et al., 2016). This finding implies that myelinated axons sprout new collaterals that are guided to the epidermis during hair growth. If the simplified arbors we previously observed represent such newly sprouted collaterals, they would not be sampled by longitudinal imaging of axons located prior to hair growth. Future time-lapse studies that survey overall innervation on each imaging day would address this discrepancy.

The degree of plasticity we observed in axons both exceeded and occurred in absence of plasticity in Merkel cells. We expected to see that changes in Merkel cells and axonal terminals occur together because they are synaptic partners, and Merkel cells are required for slowly adapting type I firing patterns (Maksimovic et al., 2014; Ikeda et al., 2014; Hoffman et al., 2018). On the other hand, previous studies have shown that touch-dome keratinocytes are responsible for gross axonal targeting to touch domes during development and maintenance in adulthood (Maricich et al., 2009; Jenkins et al., 2019; Maksimovic et al., 2014; Reed-Geaghan et al., 2016; Doucet et al., 2013). Thus, it is possible that the rate of axonal remodeling is either set intrinsically by neuronal mechanisms or is modulated by other cell types such as touch-dome keratinocytes or BMP4-expressing dermal cells, which are necessary for touch-dome innervation during development (Jenkins et al., 2019).

Although remodeling in Merkel cells is not required for axon terminal plasticity, we found that Merkel cells do alter axonal remodeling. In wild type mice, we found that contacting a Merkel cell stabilizes individual branches and is necessary for developing the kylix. Moreover, after Merkel cells are removed, branching increases throughout the arbor. Consistent with these observations, our analysis of *Atoh1* knockout mice revealed that, in the complete absence of Merkel cells, TrkC+ touch-dome axons are hyperbranched and lack kylix endings. Thus, Merkel cells stabilize axonal branches and promote ending maturation to the kylix morphology.

### Limitations of this study

As with any live-cell imaging study, one must consider the possibility that the observation itself might alter plasticity. In particular, either photodamage or skin-barrier disruption might have increased the amount of plasticity observed *in vivo*. To minimize photodamage, two-photon laser power, exposure and imaging depth was carefully controlled. We do not believe that photodamage had a significant effect on the plasticity we observed, because the degree of axonal remodeling did not correlate with total exposure. Moreover, *post hoc* comparison showed that arbors were similar in axons that had been repeatedly imaged compared with non-irradiated control axons in the same specimens.

Another potential confound is that hair removal was necessary to perform high-resolution imaging of axons in hairy skin. To minimize autofluorescence in the imaging field, fur was carefully clipped on the first day of imaging and a gentle, over-the-counter depilatory cream was used to remove hair shafts. A previous study argued that sensory neurons have intrinsic plasticity that is enhanced by fur clipping (Cheng et al., 2010). This study correlated expression of growth-related proteins and *Nefh* mRNA with days since fur-clipping but did not analyze age-matched controls; therefore, the impact of fur clipping on epithelial turnover and axonal plasticity remains unclear. Depilatory cream transiently reduces skin barrier function, which can stimulate temporary epidermal proliferation (Wahlberg, 1972). Although this is not a nervous system injury, barrier disruption could alter the dynamics of Merkel cells and terminal branches in the epidermis. Other studies have established that skin injury increases proliferation and Merkel-cell production (Tachibana & Nawa, 1980, which likely explains the higher rate of Merkel-cell addition after shaving with a razor (Wright et al., 2017). We tested two depilation protocols: one in which mice were depilated before each imaging session and one in which depilation was performed as needed to remove regrown hair. There was no difference in the amount of plasticity in Merkel cells or axon terminals when depilation groups were compared, suggesting that repeated exposure to depilatory cream did not impact plasticity. Nonetheless, the extent of injury caused by fur clipping and use of depilatory cream compared with straight-razor shaving, as well as the impact of these commonly used hair-removal techniques on skin health and neuronal plasticity, remain unclear.

### Signaling cascades that might underlie axonal plasticity

Although our results shed light on the cellular substrates of axonal plasticity, they also raise questions as to what mechanisms dictate remodeling sensory axons during epithelial homeostasis. The availability of RNA sequencing datasets for both adult Merkel cells and mature Ab sensory neurons provides an opportunity to identify candidate mechanisms (Hoffman et al., 2018; Zheng et al., 2019).

For example, it is possible that developmental mechanisms of axonal patterning remain active to control axonal remodeling in adulthood. In many nascent neurons, branch regression during development is mediated by the pro-apoptosis factors caspase and bcl-2 (Nikolaev et al., 2009; Maor-Nof & Yaron, 2013). Indeed, caspase-3, bcl-2, and bax are expressed by adult dorsal root ganglion neurons and could mediate branch removal in adulthood (Zheng et al., 2019). Additionally, BMP4 signaling is necessary for Merkel cell-neurite complex formation

during development (Jenkins et al., 2019) and altering the amount of BMP expression in the skin by either BMP or Noggin overexpression leads to changes in innervation density (Guha et al., 2004). Both touch-dome keratinocytes and a population of dermal cells beneath the axonal arbor express BMP4 in adulthood, and thus could govern axonal and Merkel-cell plasticity by modulation of BMP4 signaling (Jenkins et al., 2019). Indeed, both Aβ afferents and Merkel cells express Bmpr1a and Bmpr2, which mediate BMP4 signaling (Miyazono et al., 2010). The tangled morphology of afferents in animals lacking Merkel cells also resembled the phenotype of peripheral axons in zebrafish that harbor mutations in kinesin light chain 4; indeed, kinesin light chain 4 is expressed in sensory neurons so it is possible that this cargo transport pathway gets disrupted or overloaded since terminal branches are not able to fully mature in the absence of their target cells (Haynes et al., 2022). Furthermore, Merkel-cell afferents express the neurotrophin receptor NTrk3 (TrkC) and NT3 is necessary for postnatal maintenance of both Merkel cells and their axons (Airaksinen et al., 1996) and could govern the ongoing plasticity that we observed. Interestingly, NT3 is both highly expressed and enriched in Merkel cells compared to surrounding keratinocytes, suggesting that Merkel cells are a source of NT3 that binds to axonal NTrk3.

Molecules that govern axonal regression and growth after injury could also govern remodeling in the absence of pathophysiology. Axonal degeneration in multiple models of injury and disease depends on Sarm1, an NADase that is expressed in all neurons and results in Wallerian degeneration when activated (Osterloh et al., 2012; Gerdts et al., 2015). STAT3 and Shh play a role in new branch sprouting after axonal injury (Bareyre et al., 2011; Luo et al., 2016; Martinez et al., 2015; Berretta et al., 2016) and could be involved in regulation of ongoing plasticity in the absence of an axonal injury as both are expressed in Aβ sensory neurons. Shh in particular is a good candidate for regulation of plasticity in Merkel cell-neurite complexes as it is required in afferents to maintain Merkel cells in adulthood (Xiao et al., 2015).

Since the Merkel cell-neurite complex is a peripheral synapse, molecules that govern its maintenance could be shared with another peripheral synapse, the neuromuscular junction. Neural-derived agrin is required for post-synaptic sites on muscle fibers to differentiate and stabilize during development (Gautam et al., 1996; Jones et al., 1997). Both Agrin and synaptic Laminin are required for maintenance of neuromuscular junctions in adulthood (Samuel et al., 2012). Neurotrophin signaling via TrkB is also required for maintenance of the neuromuscular junction (Gonzales et al., 1999). Indeed, Merkel cells express some types of laminin and Aβ afferents express agrin, several types of laminin, and TrkB. Moreover, BDNF is required postnatally for slowly adapting firing patterns in response to mechanical stimulation (Carroll et al., 1998). Thus, these mechanisms that maintain the NMJ could also contribute to synapse maintenance at the Merkel cell-neurite complex.

Merkel cells express a variety of synaptic adhesion molecules, including neurexins (Haeberle et al., 2004; Hoffman et al., 2018). Unsurprisingly, sensory neurons including Aβ afferents also express many of these molecules. Some synaptic adhesion molecules, including contactin1 and Nrcam, are enriched in Aβ SA1 afferent compared with other afferent subtypes (Zheng et al., 2019). Future studies are needed to explore the expression of these molecules in nerve terminals that make contact with Merkel cells, and to determine whether they are necessary for terminals to stabilize and form kylikes.

### Conclusions

Together, our data show the remarkable level of plasticity in peripheral touch receptors during epithelial homeostasis but raise questions on how sensory stimuli are consistently encoded even though peripheral end organs are undergoing almost constant plasticity. Although Merkel cells are not required for axons to target the touch dome, they play a role in branch organization, refinement and stabilization. Moreover, our findings set the stage to identify the molecules governing peripheral axon morphology and plasticity in models of pathophysiology.

## Methods

### Animals

Animal use was conducted according to guidelines from the National Institute of Health’s Guide for the Care and Use of Laboratory Animals and was approved by the Institutional Animal Care and Use Committee of Columbia University Medical Center. Mice were maintained on a 12 h light/dark cycle, and food and water was provided *ad libitum*. Both sexes were used unless indicated otherwise. The following strains were used: Atoh1-A1GFP (RRID: IMSR_JAX:013593; MGI: *Atoh1^tm4.1Hzo^*; Rose et al., 2009), TrkC-tdTomato (RRID: IMSR_JAX:030292; MGI: *Ntrk3^tm2.1Ddg^*; Bai et al., 2015), Sarm1^+/-^ (RRID: IMSR_JAX:018069; MGI: *Sarm1^tm1Aidi^*; Kim et al., 2007), C57Bl/6J (RRID: IMSR_JAX:000664), K14^Cre^ (RRID: IMSR_JAX:018964; MGI: Tg(Krt14Cre)1Amc; Dassule et al., 2000), Atoh1^LacZ^ (RRID: IMSR_JAX:005970; MGI: Atoh1^tm2Hzo^; Ben-Arie et al., 2000), and Atoh1^flox^ (RRID: IMSR_JAX:008681; MGI: Atoh1^tm3Hzo^; Shroyer et al., 2007). Mice used in *in vivo* time-lapse imaging studies were *Sarm1^+/+^*; *Atoh1^A1GFP/A1GFP^*;*TrkC^tdTomato/+^*. We also analyzed mice lacking Merkel cells, *K14^Cre/+^; Atoh1^fl/lacz^; TrkC^tdTomato/+^*, and littermate controls, *K14^+/+^; Atoh1^fl/+^; TrkC^tdTomato/+^*. Figure 1 includes previously published data in P66 female C57Bl6/J mice (Marshall et al., 2016).

### Preparation for in vivo imaging

*In vivo* longitudinal experiments began when mice were in telogen and continued every 3 days until they had completed an entire hair growth cycle. Anesthesia was induced with 4% isoflurane in 100% oxygen. Once anesthesia was sufficiently deep, mice were moved to an imaging platform where they were placed in a bite bar with an anesthesia nose cone. Temperature was monitored and maintained at 35.5-37.5°C; anesthesia was maintained with 1–2% isoflurane in oxygen. Anesthesia and temperature were maintained using a SomnoSuite with RightTemp module (Kent Scientific).

Fur on top of the head was trimmed with electric clippers and surgical scissors on the first day of imaging (day 0). Remaining hair was depilated with Surgi-cream for 1–2 minutes; this was repeated either for each imaging session or as needed to maintain a clear imaging field as hair grew back during the hair-growth cycle (2 mice in each preparation group). Depilatory cream was cleaned from the skin with cotton balls soaked in warm water that were gently wiped across the skin until the cream was removed. Once the imaging field was clear, mice were placed in ear bars and a marker dot was made between the eyes along the midline to aid in orientation and focusing of the objective. After imaging, immersion gel was cleaned off with water; hand cream (Neutrogena Norwegian Formula Hand Cream) was applied to the depilated region. Hand cream was also applied on days between imaging sessions.

### Conventional epifluorescence microscopy and registration of arbors and hair-cycle stage

Imaging was performed with a 20X water-immersion objective (Olympus, NA=1.0). GenTeal Lubricant Eye Gel was used as an immersion medium. Conventional epifluorescence z-stacks were acquired using a CCD camera (SciCamPro, Scientifica) and Micro-Manager software (Edelstein et al., 2010). Red and green epifluorescence stacks were acquired in series with a step size of 10 µm.

Guard hairs were located on the first day of imaging. These hairs are notably larger and more autofluorescent than the other hair types in the mouse coat; additionally, these hairs are adjacent to touch domes and therefore should have TrkC-tdTomato labeling nearby. The coordinates relative to the marker dot were recorded (Scientifica MMBP Stand Alone motorized stage). We returned to the same coordinates relative to newly placed marker dots on subsequent days and then searched for guard hairs and labeled axons. Once a labeled axon was found, a conventional epifluorescence z-stack was acquired and compared with epifluorescence z-stacks from previous sessions, specifically, making sure that the guard hair and surrounding hair follicles were in the same arrangement as in the previous session. Coordinates were also used to find arbor locations relative to each other, such that once a previously recorded arbor was located it served as a reference point for finding other axons. Hair follicle positions and axon identity were also confirmed post hoc.

Conventional epifluorescence z-stacks were also used to distinguish the hair-cycle stage at each session. Actively growing hairs appeared black in epifluorescence images, whereas club hairs appeared bright. The diameter of the visible portion of the growing hair increased as the hair progressed through anagen before shrinking in catagen and returning to a bright, autofluorescent spot in telogen. Peak anagen denotes the session with the largest diameter actively growing hair (see **Figure S1**).

### Two-photon microscopy

Once axons were registered, two-photon stacks were acquired. Our microscope could readily switch between conventional epifluorescence and two-photon modes without having to move the objective, enabling acquisition of two-photon images of the locations registered with epifluorescence. Two-photon images were acquired using a resonant-scanning multiphoton system (Scientifica) and ScanImage software (Pologruto et al., 2003). The laser used for excitation was a femtosecond Ti-sapphire oscillator (Mai Tai HPDS-232, Spectra Physics) that tuned was tuned to a wavelength of 940 nm and controlled by a Pockels Cell (Conoptics). GFP and tdTomato fluorescence were simultaneously acquired. TdTomato fluorescence was collected with a GaAsP photomultiplier tube (Scientifica, S-MDU-PMT-50-65) and green fluorescence was collected with a raw PMT (Scientifica, 2PIMS-PMT-45 Red Raw).

Image zoom was adjusted to maximize the number of pixels capturing the arbor while attempting to eliminate as much of the hair-follicle autofluorescence as possible. Sometimes the hair follicle could not be completely excluded from the frame and the Scan Image Power Box utility was used to set the laser power over the hair follicle to 0. Black anagen hairs obscured two-photon images during some anagen sessions; these were excluded from analysis (see **Table S2**). The bounds of z-stacks were set to include the whole afferent arbor and the stack was acquired at 1 µm step size. One hundred and fifty frames were averaged per plane. For publication, representative images were cropped to regions of interest and output levels were linearly adjusted in photoshop to ensure the histogram filled the dynamic range. Gamma was not changed.

### Whole-mount immunohistochemistry, tissue clearing, and confocal microscopy

For time-lapse imaging mice, skin from the longitudinal imaging site was dissected immediately after the conclusion of *in vivo* imaging on the last day of the longitudinal experiment. Other skin samples were collected from the back for additional immunostaining. Body sites that were not already depilated from longitudinal *in vivo* studies were depilated (Surgi-cream), then skin samples were dissected. Care was taken to cut the skin of the head to allow re-orientation for confocal registration of afferents imaged *in vivo*.

Whole-mount immunohistochemistry was performed as previously described (Marshall et al., 2016). Briefly, tissue was fixed in 4% paraformaldehyde (PFA) overnight, washing in 0.03% triton-X PBS (PBST) and incubated in primary antibody for 5–7 days at 4°C. Primary antibodies used were as follows: rat anti-K8 (Developmental Studies Hybridoma Bank, TROMA-I; RRID: AB_531826), chicken anti-NFH (Abcam, ab4680; RRID: AB_304560), rabbit anti-NFH (Abcam, ab8135; RRID: AB_306298), rabbit anti-DsRed (Clontech, 632496; RRID: AB_10013483), and chicken anti-GFP (Abcam, ab13970; RRID: AB_300798). After 10–16 h of washes in PBST, samples were incubated for 48 h at 4°C in secondary antibodies: goat Alexa Fluor-conjugated antibodies (Invitrogen) directed against rat (Alexa Fluor 647, A21247; Alexa Fluor 594, A11007), chicken (Alexa Fluor 488, A32931; Alexa Fluor 594, A11042), or rabbit (Alexa Fluor 594, A11012; Alexa Fluor 647, A21244) IgG. After labeling, tissue was dehydrated progressively in 25–100% methanol and cleared using a 2:1 benzyl benzoate/benzyl alcohol solution.

*Sarm1^+/+^; Atoh1^A1GFP/A1GFP^; TrkC^tdTomato/+^* specimens were imaged in three dimensions on a Nikon Ti Eclipse confocal microscope equipped with 20X, 0.75 NA objective lens in the Confocal and Specialized Microscopy Shared Resource of the Herbert Irving Comprehensive Cancer Center at Columbia University, supported by NIH grant P20CA013696. C57Bl/6J data in **Figure 1** and Atoh1^CKO^ data in **Figure 7** were imaged on a Zeiss Exciter confocal microscope with a 20X, 0.8 NA or 40X, 1.3 NA objective lens, respectively; these experiments were conducted in the Columbia University EpiCure Center (NIAMS P30AR044535).

Quantification was performed in unprocessed axial stacks. For publication, representative images were cropped to regions of interest and output levels were linearly adjusted in photoshop to ensure the histogram filled the dynamic range. Gamma was not changed.

### Analysis of two-photon and epifluorescence images

Epifluorescence images were used to register arbors across sessions. To confirm that hair follicles were in the same pattern, single z planes were exported from ImageJ to Adobe Illustrator, where hair follicle locations were noted and confirmed to be in the same arrangement around the arbor across sessions. Arbors were included in plasticity analyses only if they had the same hair-follicle arrangement in every session. Axons and Merkel-cell clusters were also excluded from analysis if they were imaged in fewer than 60% of sessions (**Table S2**). A total of 20 arbors from 4 mice passed these criteria and were analyzed.

### Measurement of Merkel-cell contact areas

To determine the size threshold between kylikes and boutons, we measured the length and width of each ending at each timepoint in ImageJ. Ending area was approximated using the formula for ellipse area:

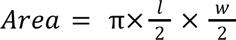

The histogram of ending ellipse areas was fitted with a bimodal gaussian distribution in Prism (Graphpad) and the threshold between large kylix endings and small bouton endings was defined as the crossing point of the two component gaussians that when summed together made the bimodal gaussian.

### Plasticity analysis

Two-photon z-stacks were motion corrected (SIMA, Kaifosh et al., 2014). The following parameters were manually tracked from session to session: Merkel cell addition, removal, and movement, terminal branch addition, removal, growth, and regression, as well as stability of both Merkel cells and branches. Merkel cells were defined as the same cell as the previous session if they shared either cell shape or location with a Merkel cell in the previous session; if both cell shape and position were different from existing cells, that cell was determined to be a new cell. Merkel cells could only be counted as “moved” if their shape was the same as a cell imaged at the previous timepoint and a line could be drawn from the current position to the previous position without going through any other Merkel cells. Additionally, the quality of the contact between Merkel cells and terminal branches was tracked from session to session. If the same branch changed from a kylix to a bouton, it was scored as a regression event, reflecting the decrease in ending area; similarly, if a terminal changed from a bouton to a kylix, it was scored as a growth event. One axon was not able to be fully analyzed using these methods because its branching structure was too complex to confidently register terminals across sessions (**Table S2**; **Figure S2**). However, Merkel-cell counts and their contact types were included in analyses. All parameters that report changes across sessions were normalized to the number of imaging days that occurred between the sessions being compared (i.e. changes from days 6–9 would be normalized to 1 session as they were consecutive imaging sessions, but if day 6 was compared to day 12 changes would be normalized to 2 sessions).

### Data Analysis and Statistics

Statistical analyses (one- and two-way ANOVA, t tests (paired, unpaired, parametric, nonparametric), linear regression, 2×2 Fisher’s exact test, survival curve analysis, curve fitting) were performed with Prism 9 (Graphpad). For Fisher’s exact tests of tables that were larger than 2×2, Vassar Stats online Fisher’s exact calculator was used. For parametric data with three or more groups, one-way ANOVAs were followed by Tukey’s post hoc analysis for between-group comparisons. Student’s two-tailed *t* test was used to compare means of two normally distributed groups. Categorical data were compared using two-tailed Fisher’s exact test. Two-way ANOVA with Tukey’s post hoc was used to compare the number of Merkel cells and neurites in the first and last session. Custom MATLAB scripts were used to tally numbers of Merkel cells and terminals fitting into groups included in analyses (i.e. the number of terminals that showed plasticity), as well as compute plasticity parameters.

### Plasticity parameters

Plasticity of Merkel cells and terminal branches was summarized by a plasticity index, which is defined for Merkel cells and terminal branches, respectively as follows:

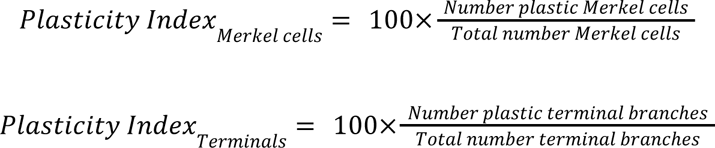

These plasticity indices encompass all plasticity events (for Merkel cells, addition, removal, movement; for terminal branches, addition, removal, growth, regression, change in ending morphology) such that an index of 0 represents stability of that cellular component, either Merkel cells or terminal branches, of the arbor and 100 represents that all Merkel cells or branches were changed. This index is calculated on a session to session basis, meaning that only changes since the last session are included.

In addition to overall plasticity, we also calculated Merkel cell removals and a regression index for terminal branches. Merkel-cell removals was simply defined as the number of Merkel cells removed since the last session. We first totaled the number of terminals (*Nt*) going through any kind of regression:

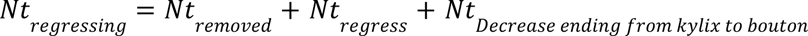

Then, converted to a percentage by dividing by the number of plastic terminal branches and multiplying by 100.

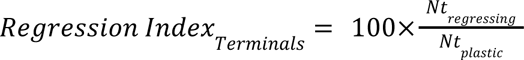

### Permutation analysis

Lists containing the temporal ordered plasticity parameters were manually aligned according to external factors, including two-photon session number and hair cycle. Using a custom MATLAB script, the temporal profiles of all plasticity parameters for each touch dome were shuffled 1,000 times. Pairwise correlation matrices were computed; the median correlation coefficient was measured for each of the shuffled cases. To determine the *P* value for whether the median correlation coefficient of the true data was different from the shuffled data, we counted the proportion of shuffled cases with medians greater than the experimental median:

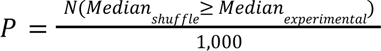

The cumulative frequency of each shuffled case was computed and averaged to obtain the average±SD across all shuffled instances.

## Acknowledgements

We thank Drs. Benjamin Hoffman, Georgia Pierce, Dan Kato, and Sam Benezra for assistance with establishing the *in vivo* imaging preparation. Dr. David Ginty assisted with TrkC-tdTomato mice. Drs. Wesley Grueber, Carol Mason, Ulrich Hengst, Randy Bruno, A. James Hudspeth, Yalda Moayedi, Diana Bautista and members of the Lumpkin, Grueber, and Bautista laboratories provided helpful discussions. Imaging core facilities were supported by the Columbia University EpiCURE Center (NIAMS P30AR069632) and the Confocal and Specialized Microscopy Shared Resource of the Herbert Irving Comprehensive Cancer Center (NCI P30CA013696). Figure 2A was created with BioRender.com. RCC was supported by NINDS F31NS103439, NINDS T32NS064928, and NICHD T32HD007430. BAJ was supported by NINDS F31NS094023. This research was funded by NINDS R01NS073119 and NIAMS R01051219 (to EAL).

## Author Contributions

**Conceptualization:** RCC, EAL

**Methodology:** RCC, BAJ

**Software:** RCC

**Formal analysis:** RCC, EAL

**Investigation:** RCC, BAJ

**Data curation:** RCC, EAL

**Writing—original draft:** RCC

**Writing—review & editing:** RCC, EAL

**Visualization:** RCC

**Supervision:** EAL

**Funding acquisition:** RCC, BAJ, EAL

---

**Figure S1, relevant to Figure 2.**
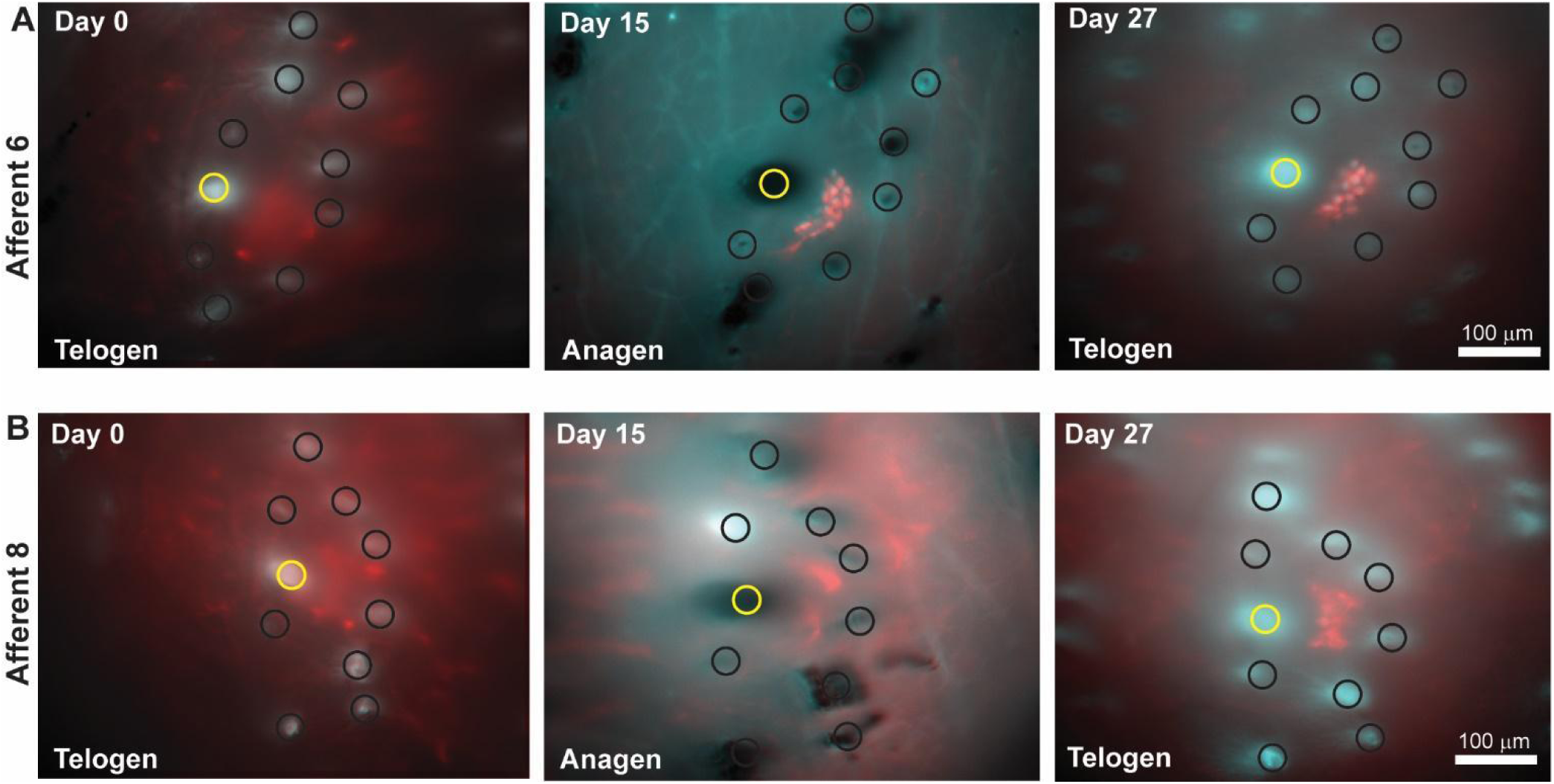
Hair follicles are unique landmarks to register afferent locations for several weeks. Adult mice go through hair growth cycles after post-natal hair morphogenesis. The first cycle usually begins between P28 and P35 in C57Bl6/J mice, but differs with genetic background, sex, body size, and stress level. Telogen, the resting phase of hair growth, has small hair follicles with hairs that are not actively growing. Anagen, the phase of active hair follicle growth, is a period of massive cellular proliferation. The skin thickens to accommodate larger hair follicles that are actively growing hair and appears back from the long lengths of actively growing hair that are present under the skin. Cells throughout the skin turnover to expand the thickness of the skin to accommodate enlarged hair follicles. The final phase, catagen, is a short phase in which follicles actively regress back to their resting size. Epifluorescence z-stacks were used to register the locations of hair follicles surrounding touch domes. Hair follicle locations were used to confirm the identity of each afferent that was part of the time-lapse imaging study in an un-biased manner. **A–B**) Single z planes of epifluorescence z-stacks of the same areas taken throughout time-lapse imaging. Yellow circles indicate guard hair follicle immediately adjacent to touch dome. Black circles indicate surrounding hair follicles used as landmarks. **A** and **B** are different touch domes from the same animal; afferent ID indicated on left of each row. Imaging day indicated in top left corner; hair-cycle stage indicated in bottom left corner. Note that the relative positions of hair follicles surrounding individual touch domes are unique and persist throughout hair cycle stages for ∼1 month. Cyan: Atoh1-GFP; red: TrkC-tdTomato.

**Figure S2, relevant to Figure 2.**
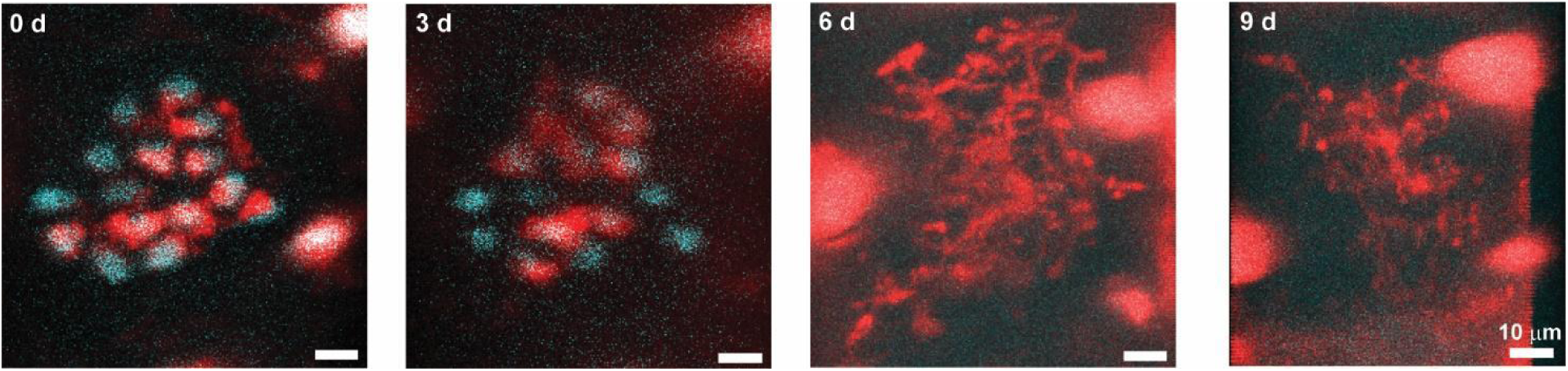
An afferent that had complete loss of the Merkel-cell cluster showed complex branching structure that could not be quantified with existing methods. *In vivo* time series of touch dome no. 1 from mouse M2. Individual branches could not be registered with confidence after day 3. Note hyper-branched morphology on day 6 that started to refine on day 9. Cyan: Atoh1-GFP; red: TrkC-tdTomato.

**Figure S3, relevant to Figure 2.**
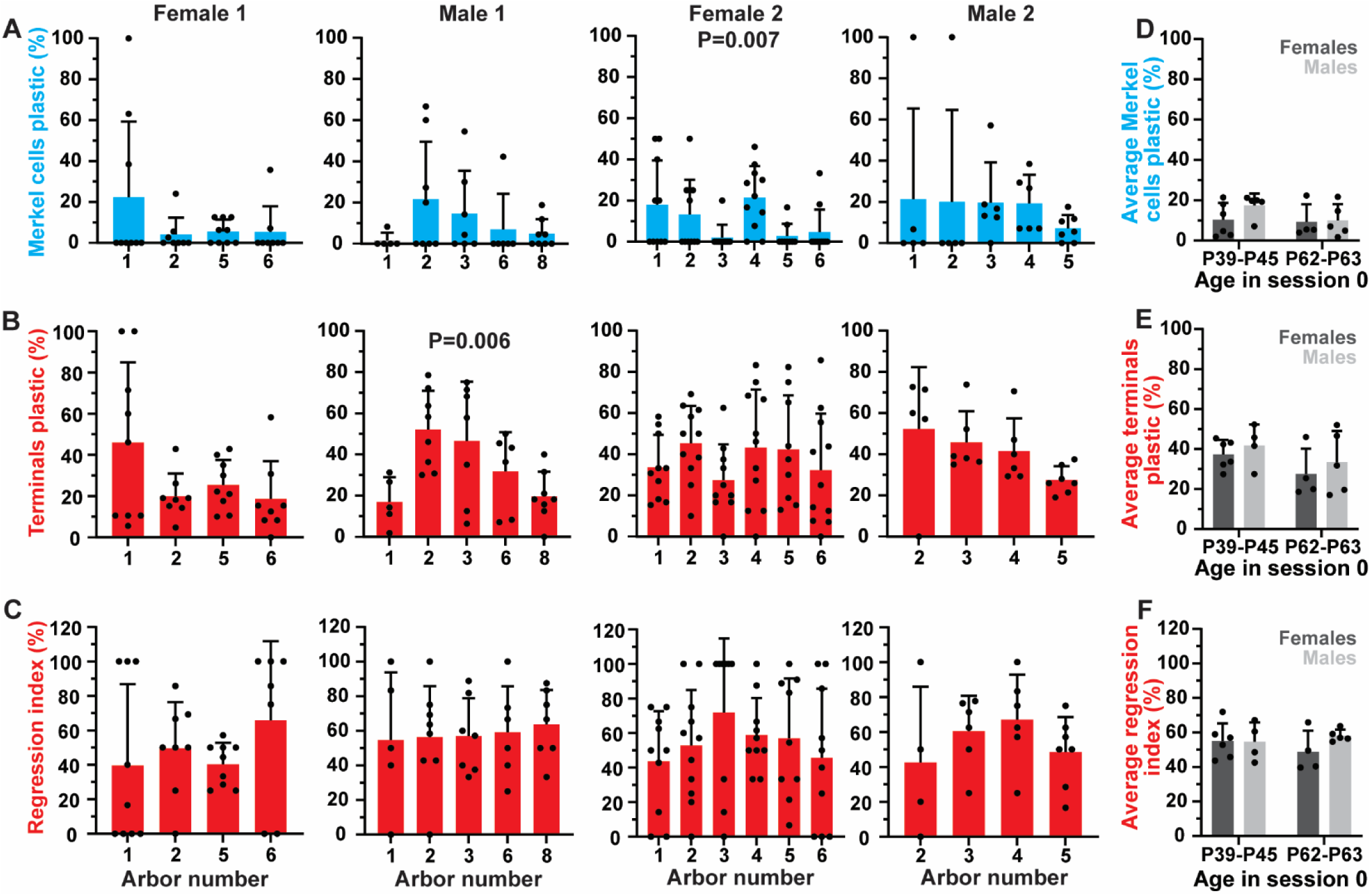
The percentage of Merkel cells and axon terminals that showed plasticity, as well as regression index, is heterogeneous across touch domes within individual mice but does not differ with sex or age. (**A–C**) Comparison of the range of plasticity in Merkel cells (**A**) and terminal neurites (**B**), as well as regression index (**C**) between imaging sessions for each touch dome analyzed. Note the variance in the amount of plasticity across afferents in the same animal. Bars indicate means, error bars indicate SDs, and dots show all time points. P values are results from Kruskal-Wallis nonparametric ANOVA (F2 Merkel cells) or ordinary one-way ANOVA (M1 terminals). (**D–F**) Comparison of average plasticity in Merkel cells (**D**) and terminals (**E**), and regression index (**F**) separated by age and sex. No significant interaction of either age or sex by two-way ANOVA for any parameter.

**Figure S4, relevant to Figure 5.**
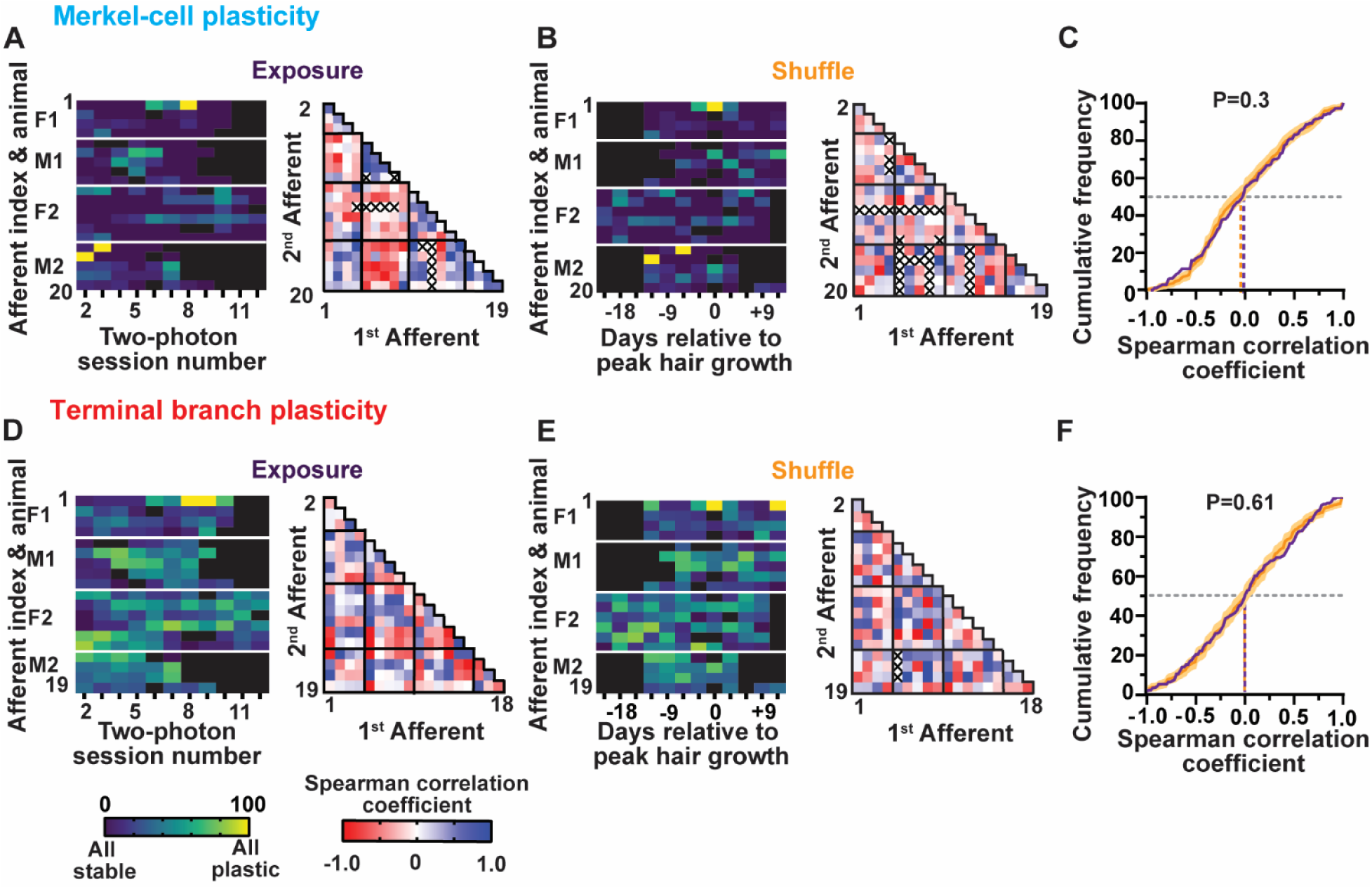
Two-photon laser exposure does not synchronize Merkel-cell or axon plasticity across arbors. Analysis of Merkel-cell plasticity index (**A–C**) and terminal branch plasticity index (**D–F**) for every afferent used in analysis following the same organization as **Figure 5**. (**C,F**) Cumulative histogram of correlation coefficients from exposure-aligned experimental data and mean±SD cumulative histogram from correlation matrices of shuffled permutations. *P* values: permutation test. Gray dashed line indicates 50%, colored dashed lines indicate median correlation coefficients for each group. White lines in **A, B, D, E** indicate the arbors from each animal; black squares indicate days with unanalyzable or missing data.

**Figure S5, relevant to Figure 7.**
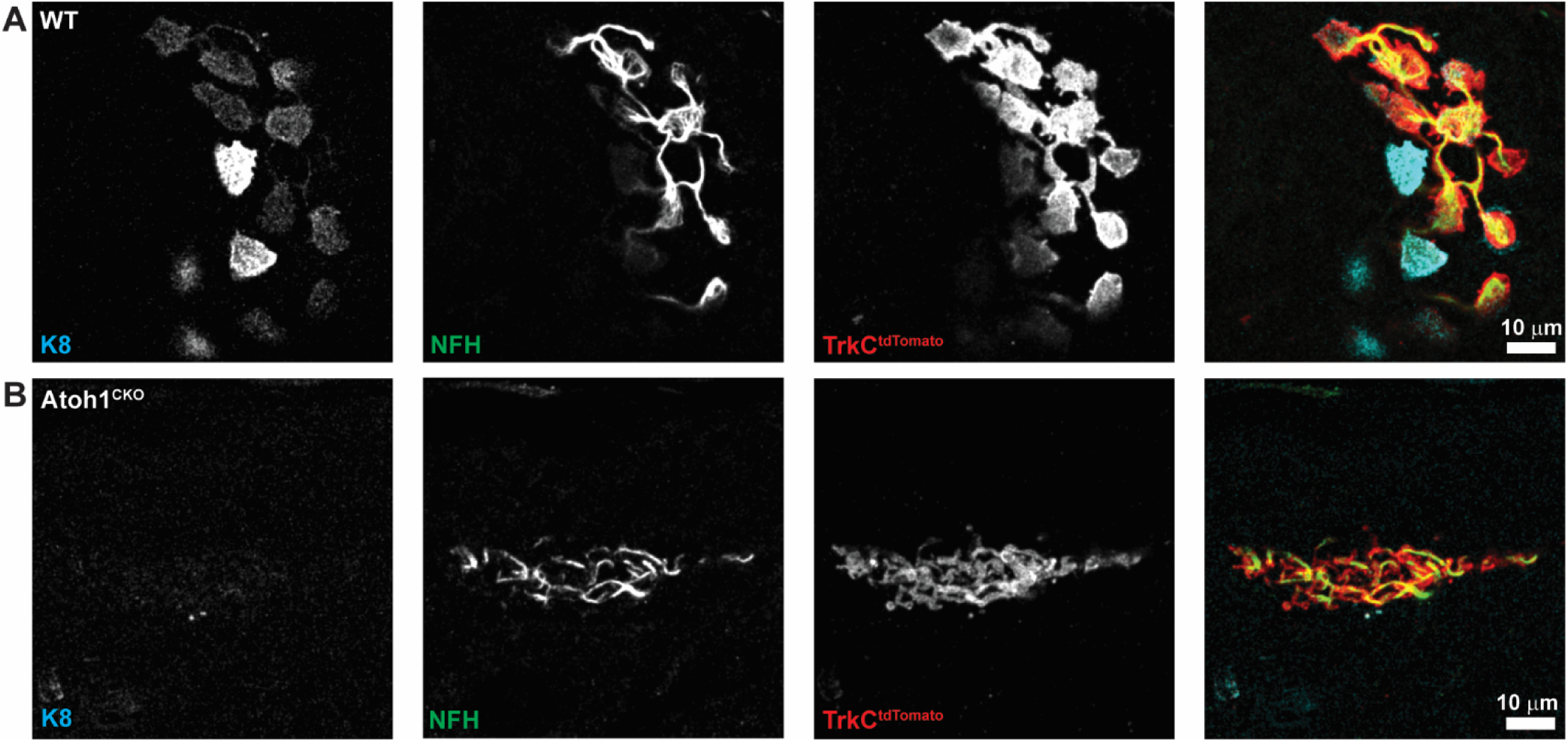
Kylix formation requires Merkel cells. (**A–B**) Axial projections from substacks of wildtype (**A**) and Atoh1^CKO^ (**B**) touch domes from **Figure 7**. Projections are of 3 z planes to show the tangled branches of the Atoh1^CKO^ animal and absence of mature kylikes. K8 (cyan), NFH (green) and DsRed to amplify TrkC-tdTomato (red).

**Table 1.**
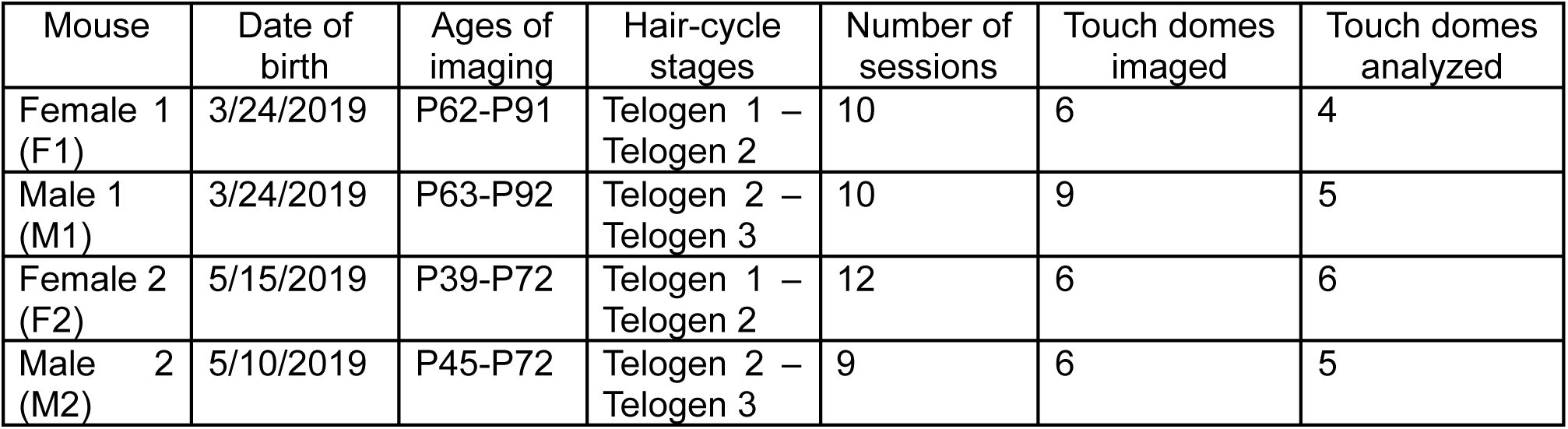
Information for individual mice.

**Table 2.** All recorded touch domes and whether they were included in analysis.

| Animal | Touch dome number | Fraction of sessions imaged | Included? | Peak anagen session included? |
| --- | --- | --- | --- | --- |
| F1 | 1 | 10/10 | Yes | Yes |
| F1 | 2 | 10/10 | Yes | No |
| F1 | 3 | 4/10 | No | N/A |
| F1 | 4 | 4/10 | No | N/A |
| F1 | 5 | 10/10 | Yes | Yes |
| F1 | 6 | 9/10 | Yes | No** |
| M1 | 1 | 8/10 | Yes | No |
| M1 | 2 | 9/10 | Yes | Yes |
| M1 | 3 | 8/10 | Yes | Yes |
| M1 | 4 | 1/10 | No | N/A |
| M1 | 5 | 1/10 | No | N/A |
| M1 | 6 | 8/10 | Yes | No |
| M1 | 7 | 1/10 | No | N/A |
| M1 | 8 | 10/10 | Yes | Yes |
| M1 | 9 | 1/10 | No | N/A |
| F2 | 1 | 12/12 | Yes | Yes |
| F2 | 2 | 12/12 | Yes | Yes |
| F2 | 3 | 11/12 | Yes | No** |
| F2 | 4 | 12/12 | Yes | Yes |
| F2 | 5 | 12/12 | Yes | No |
| F2 | 6 | 12/12 | Yes | Yes |
| M2 | 1 | 6/9 | Yes* | No* |
| M2 | 2 | 7/9 | Yes | Yes |
| M2 | 3 | 7/9 | Yes | Yes |
| M2 | 4 | 7/9 | Yes | Yes |
| M2 | 5 | 9/9 | Yes | No |
| M2 | 6 | 5/9 | No | N/A |

*\*Afferent whose terminal branches could not be registered with confidence and was only included in analyses of Merkel cells and contacts.*
*\*\*Afferents that were not found and thus not imaged during anagen.*

